# Neuromodulatory Control of Cortical Function: Cell-Type Specific Regulation of Neuronal Information Transfer

**DOI:** 10.64898/2026.03.13.711516

**Authors:** Nishant Joshi, Xuan Yan, Niccolo Calcini, Payam Safavi, Asli Ak, Koen Kole, Sven van der Burg, Tansu Celikel, Fleur Zeldenrust

## Abstract

Neuromodulatory systems such as dopamine and acetylcholine enable cortical circuits to rapidly shift between behavioral states, yet it remains unclear how receptor-specific signaling reshapes what single neurons compute. Here we combined whole-cell recordings from excitatory and inhibitory neurons in layer 2/3 of mouse somatosensory cortex with information-theoretic analyses under frozen-noise stimulation. We quantified stimulus-to-spike information transfer and profiled each neuron across four functional domains: passive membrane biophysics, action potential dynamics, intrinsic adaptation currents, and input feature selectivity captured by spike-triggered averages. Activation of D1, D2, or M1 receptors produced cell-type and receptor-specific changes in fractional information and firing output, demonstrating that neuromodulators regulate encoding beyond simple gain control. Unsupervised clustering revealed that receptor activation reshapes neuronal functional identities, consistent with a dynamic computational phenotype. To identify the organizing principle underlying these transitions, we applied multi-set correlation and factor analysis and found that neuromodulation systematically reorganizes covariance among functional domains. In excitatory neurons, dopaminergic activation led to a decreased coordination between input feature selectivity and other functional properties while strengthening coordination among response based features such as spike dynamics and adaptation. Inhibitory neurons, by contrast, generally exhibited increased coordination across domains. These findings identify neuromodulation as a reconfiguration signal that reshapes not only individual cellular properties but also the architecture linking them, thereby dynamically expanding the computational repertoire of cortical circuits.

## Introduction

Neuromodulators such as dopamine and acetylcholine are essential for flexible brain function, supporting state-dependent changes in attention, learning, memory, and sleep (1–3). Rather than directly driving spiking, these signals tune how neurons transform synaptic input into output (4–9), thereby altering circuit computation without changing the underlying wiring diagram. Disruptions of neuromodulatory pathways are central to neurological and psychiatric disorders, including Parkinson’s disease and schizophrenia, underscoring the need for mechanistic principles that link receptor-level effects to computation (10–12). Yet most work has characterized neuromodulation using scalar descriptors such as firing rate, gain, or spike-frequency adaptation (5, 13, 14), which capture important outcomes but do not directly quantify what neurons encode or how reliably they transmit information. A central open question is therefore how neuromodulators reorganize information processing across cortical populations, and whether these effects differ across cell types.

A major challenge in understanding the effect of neuromodulators on information processing is that neuronal computation is distributed across interacting functional domains. Passive membrane properties shape filtering and excitability; action potential dynamics constrain output timing and precision; adaptation currents regulate temporal responsiveness, and stimulus selectivity reflects the joint influence of these intrinsic features on input integration as well as effects of the network. Neuromodulation act not only by adjusting average single properties in a group of cells, but also by reorganizing different cellular properties (15–19) in an organized manner, redefining the “rules” by which cellular mechanisms combine to generate stimulus-driven spiking. We therefore asked whether receptor-specific neuromodulation reconfigures population-level relationships among neuronal properties of cortical neurons by altering both (i) stimulus-to-spike information transfer and (ii) the covariance structure linking key functional domains, and whether these effects differ between excitatory and inhibitory cells. To address this, we utilized frozen-noise stimulation (20) in whole-cell recordings from neurons in layer 2/3 mouse somatosensory cortex, systematic feature extraction across four domains, and multivariate analyses that distinguish coordinated from independent modulation of neuronal function.

We quantified how neuromodulation affects the amount of information a neuron transmits. First, we performed singleunit patch clamp recordings using a frozen noise stimulus and using an information-theoretic analysis (20), to estimate the fractional information (FI), the fraction of input entropy captured by the spike train, for each neuron during control and agonist conditions. We hypothesized that receptor activation might change information transfer in a cell-typespecific manner. Second, we analyzed the effects of neuro-modulation in a high-dimensional electrophysiological feature space. From each recording, we extracted four sets of functional attributes that capture different aspects of neuronal function (21): (1) action potential (AP) waveform and firing dynamics, (2) passive biophysical (PB) properties of the membrane, (3) adaptation currents (AC) describing intrinsic spike-frequency adaptation, and (4) the spike-triggered average (STA) of the input current (reflecting input feature selectivity). These feature sets range from passive properties to active, input-driven characteristics. We then examined whether receptor activation reorganized the functional identity of neurons. Specifically, we asked if neurons that clustered together using an unsupervised method in high-dimensional space during control conditions would still cluster together during neuromodulatory receptor activation. Finally, we applied Multi-set Correlation and Factor Analysis (MCFA) (22) to assess whether neuromodulation alters the correlations between attribute sets, reflecting a global reconfiguration of each neuron’s computational architecture. Together, we provide an openly available electrophysiological dataset to study functional changes in individual neurons during neuromodulation, including raw recordings, extracted feature sets, and information-transfer measures. Using this resource, our analyses reveal how dopaminergic and cholinergic modulation reshape the functional landscape of cortical neurons. Neuromodulators thus emerge not merely as simple “gain” modulators, but as architects of neural computation that reassign the relationships between neuronal properties in a cell-type- and receptor-specific fashion.

## Results

### Neuromodulation selectively alters information transfer in excitatory and inhibitory neurons

In order to understand the effect of neuromodulation on information transfer, we first ask whether neuromodulation shifts information encoding, measured as Fractional Information (FI, see supplementary Methods), at the population level, and then assess how strongly individual neurons change relative to their own baseline and whether these changes depend on cell type. For population comparison, we calculated FI for individual neurons during control and agonist conditions (see Methods) and compared the population distributions of FI values between control and agonist conditions (Wilcoxon rank-sum tests) for each cell-type. For population comparison, we first established a baseline trial-to-trial variability by comparing the distribution between repeated control trials. We found that although the inhibitory population showed no significant change, the excitatory population showed a significant decrease (*p <* 0.001, Table S4, Fig 1A) in FI for the second trial.

**Fig. 1.**
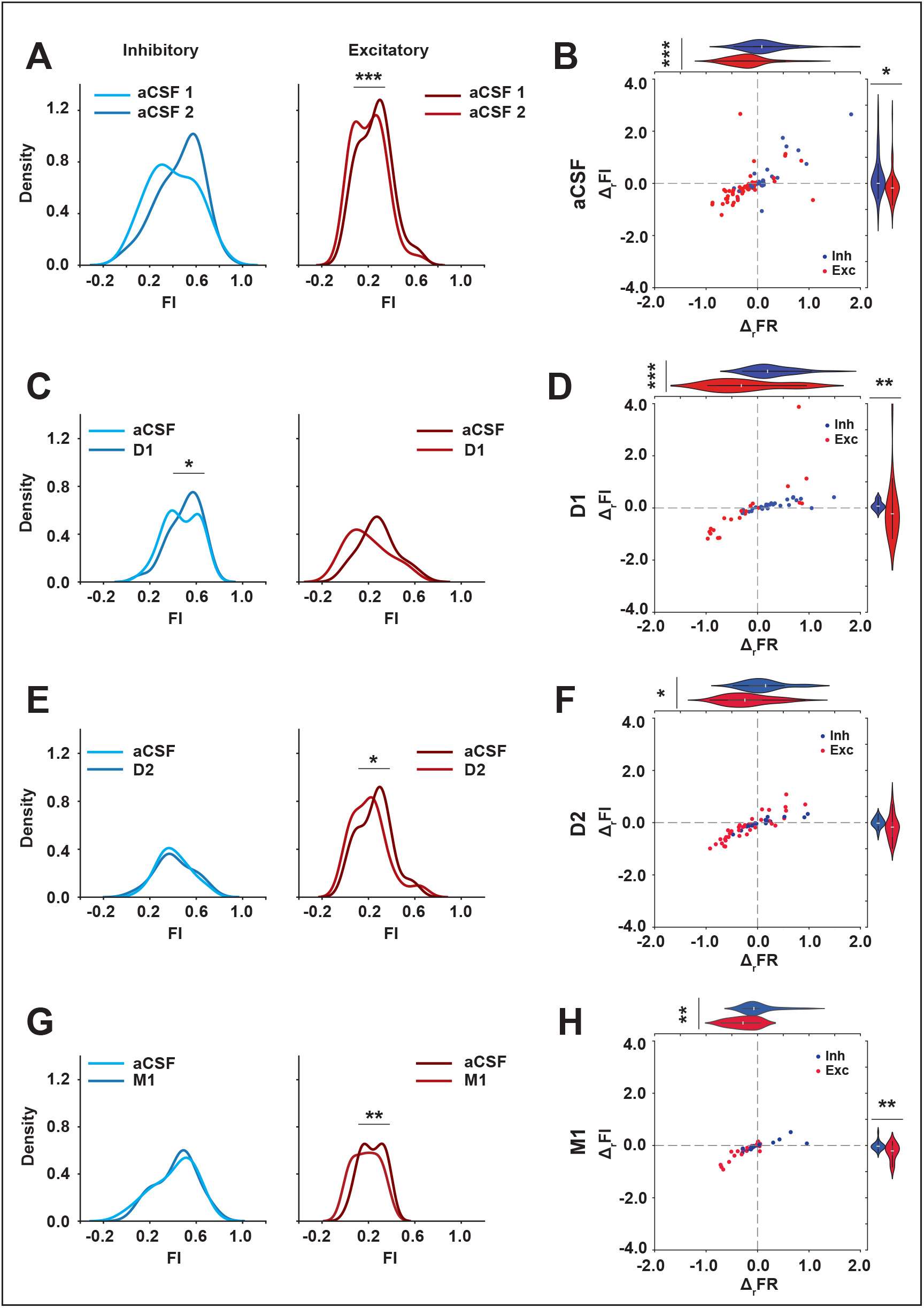
Neuromodulation changes the amount of information transferred about the stimulus in a cell specific manner. (**A**) Distribution of measured FI for repeated control trial of the same neuron for excitatory and inhibitory neurons along with control trials. (**B**) Normalized relative change in firing rate and information transfer between repeated control trials of the same neuron in a excitatory (red) and inhibitory (blue) neurons. It can be seen that the distribution of relative change in firing rates and transferred information is significantly different for excitatory neurons (red) compared to inhibitory neurons (blue). (**C**) Distribution of measured FI for control and D1 conditions respectively for excitatory and inhibitory neurons along with control trials. (**D**) Normalized change in firing rate (horizontal-axis) vs fractional information (vertical-axis) as a result of D1 receptor activation, compared across E (red) and I (blue) neurons. (**E**) Distribution of measured FI for control and D2 conditions respectively for excitatory and inhibitory neurons along with control trials. (**F**) Normalized change in firing rate vs fractional information as a result of D2 receptor activation for E (blue) and I (red) neurons. (**G**) Distribution of measured FI for control and M1 conditions respectively for excitatory and inhibitory neurons along with control trials. (**H**) Normalized relative change in firing rate vs fractional information as a result of M1 receptor activation compared across cell types. * *p* ≤0.05, ** *p* ≤ 0.01, *** *p* ≤ 0.001.

After establishing the baseline comparison, we compared the population distributions of FI between control and agonist trials. We found that during D1-R activation, the distribution of FI among inhibitory neurons increased significantly compared to control (*p <* 0.02), whereas the excitatory FI distribution did not show a significant shift (Figure 1C, Table S4). D2-R activation resulted in a decreased mean FI of excitatory neurons compared to control (*p≈* 0.01), but this shift was of similar magnitude to the baseline repeated control-trials variability (Figure 1A,E, Table S4), making it difficult to ascribe significance. The inhibitory FI distribution during D2-R activation remained essentially unchanged (*p≈* 0.77). M1-R activation caused a pronounced decrease in the mean FI of excitatory neurons (*p≈* 0.002), far beyond any baseline drift, while inhibitory neurons again showed no significant distribution change (Figure 1A,G, Table S4). Except in the D1-R condition, the inhibitory population’s FI distribution was remarkably stable across control and neuromodulator conditions (no significant differences for D2, or M1, see Table S4). In contrast, the excitatory population showed significant encoding losses for D1 and M1 receptor activation.

At the population level, these results indicate that neuro-modulation redistributes transferred information across cell-types, with excitatory neurons exhibiting a significantly larger changes in FI compared to control than inhibitory neurons for D2-R and M1-R activation. Notably, D1-R activation was associated with an increase in average FI among inhibitory neurons, along with a decrease in excitatory FI that did not reach significance at the population level. While population-level comparisons reveal whether neuromodulation shifts encoding across excitatory or inhibitory ensembles as a whole, they do not indicate how encoding changes are distributed across individual neurons within each population. We therefore next examined neuromodulatory effects at the single-cell level, focusing on relative changes in FI and firing rate within each neuron.

In order to assess how neuromodulation alters single-neuron encoding relative to baseline within excitatory and inhibitory populations, we first quantified how activating specific neuromodulatory receptors affects a neuron’s ability to transmit information, as well as its firing output. Each neuron’s fractional information (FI) (see Methods) and firing rate (FR) were measured for control (aCSF) and agonist conditions, and we calculated the relative changes Δ_*r*_*FI*_*ago*_ and Δ_*r*_*FR*_*ago*_ for each neuron (agonist minus control, normalized by control, denoted by Δ_*r*_; see Supplementary Methods). We compared these changes between excitatory and inhibitory populations to determine if neuromodulation affects the two cell types differently. To account for intrinsic variability, we established a baseline by analyzing two control trials (repeated control) for a subset of cells. A significant difference between excitatory and inhibitory Δ_*r*_*FI* does not imply that both cell types change significantly relative to baseline, but rather that neuromodulation affects the two populations differently at the level of individual neurons.

In the baseline condition (comparing two control trials), excitatory neurons showed a small but significant drift (Figure 1B, Table S5) even without drug: on average the change in the distribution of relative firing rate and FI is stronger in excitatory neurons (mean Δ_*r*_*FI*_*aCSF*_ *<* 0, *p≈* .01; mean Δ_*r*_*FR*_*aCSF*_ *<* 0, *p <* 0.0001) compared to inhibitory neurons. These baseline differences, although small (Cohen’s d Δ*FI*: -0.60, Cohen’s d Δ*FR*: -0.92), define the level of trial-to-trial variability in encoding against which drug effects should be interpreted.

Following activation of D1-R receptors, we observed a pronounced, cell-type-specific effect on encoding (Figure 1D, Table S5). Excitatory neurons exhibited a significant difference in FI and FR compared to inhibitory neurons at the level of single-neuron changes, despite the absence of a significant population-level shift (mean Δ_*r*_*FI*_*ago*_ and Δ_*r*_*FR*_*ago*_ for excitatory neurons were negative and significantly different from inhibitory; *p <* 0.005 for both, Figure 1D). These D1-driven relative changes in excitatory encoding far exceeded the baseline control variability (Cohen’s d Δ*FI*: -0.94, Cohen’s d Δ*FR*: -1.09, Table S5), indicating a robust effect of D1 modulation on excitatory neurons’ input-output efficacy (Figure 1D, Table S5). D2 receptor activation had more modest and mixed effects: on average, D2 activation led to a slight decrease in excitatory FI and FR, but these changes were smaller and not consistently significant when compared to inhibitory cells. Inhibitory neurons during D2-R activation remained relatively stable, with perhaps a very minor change in information transfer that did not rise above baseline variability (Figure 1F, Table S5). Muscarinic M1 receptor activation produced the greatest average changes of all: M1 agonist caused a pronounced difference in FI and FR relative to control between excitatory and inhibitory neurons (*p <* 0.005, see Table S5). These changes during M1-R activation also greatly exceeded baseline (Cohen’s d Δ*FI*: -0.95, Cohen’s d Δ*FR*: -1.04) underscoring a robust receptor-specific effect. In summary, D1-R and M1-R activation produced significantly different changes in information encoding and firing output between excitatory and inhibitory neurons. D2-R activation yielded weaker or more variable cell-type differences. These findings indicate that neuromodulatory influences on stimulus encoding are not uniform, but depend on both receptor type and neuronal identity. These effects were strongly noticeable during D1-R and M1-R receptor activation compared to D2-R activation.

### Receptor-specific modulation reshapes functional neuronal classification

After showing the effects of neuromodulation on cellular information transfer, we ask whether it alters the functional identity of neurons as defined by high-dimensional electrophysiological features. Neuronal cell types are traditionally classified by intrinsic features such as action potential shape and passive membrane properties, which can distinguish subtypes of excitatory or inhibitory cells (23, 24).

Recent work shows that a neuron’s classification can be depend on the input context (21, 25, 26). Given our findings above, that neuromodulation changes the amount of information different cells transmit, we hypothesize that neuromodulators might also reconfigure the relationships among a neuron’s intrinsic and input-driven attributes, effectively changing how neurons cluster in functional feature space.

To investigate the aforementioned hypothesis further, we performed an unsupervised clustering of neurons using sets of functional attributes (see Methods), namely Action Potential (AP), Passive Biophysical (PB), Adaptation Current (AC) and Spike-triggered Average (STA). We compared the resulting clusters during control versus agonist conditions (Figure 2, S2). Before analyzing drug effects, we first verified that our clustering approach was reliable and not overly sensitive to experimental variability. Specifically, for a subset of neurons with two control trials, we clustered the cells using their attributes from the first control trial and compared the resulting clusters with those obtained from the second control trial. We found a high degree of correspondence between cluster assignments from trial 1 vs. trial 2 for many attributes, particularly for waveform and passive membrane features, indicating that these functional classifications were stable across repeated control measurements (Figure S3). Attributes such as AP, AC, and STA showed a somewhat lower correspondence between control trials, likely reflecting inherent trial-to-trial variability in these measures. We also visualized the overlap of the two control trials and observed substantial alignment between the embedding generated using a UMAP technique (Figure S3), further suggesting that any differences in clustering due to neuromodulation that we will report on are not simply artifacts of recording noise or drift. We tested the sensitivity of the clusters to hyperparameters and chose the clusters with the highest modularity index score for each condition and trial, which indicates cluster consolidation (see Figure S6, S7 & S8).

**Fig. 2.**
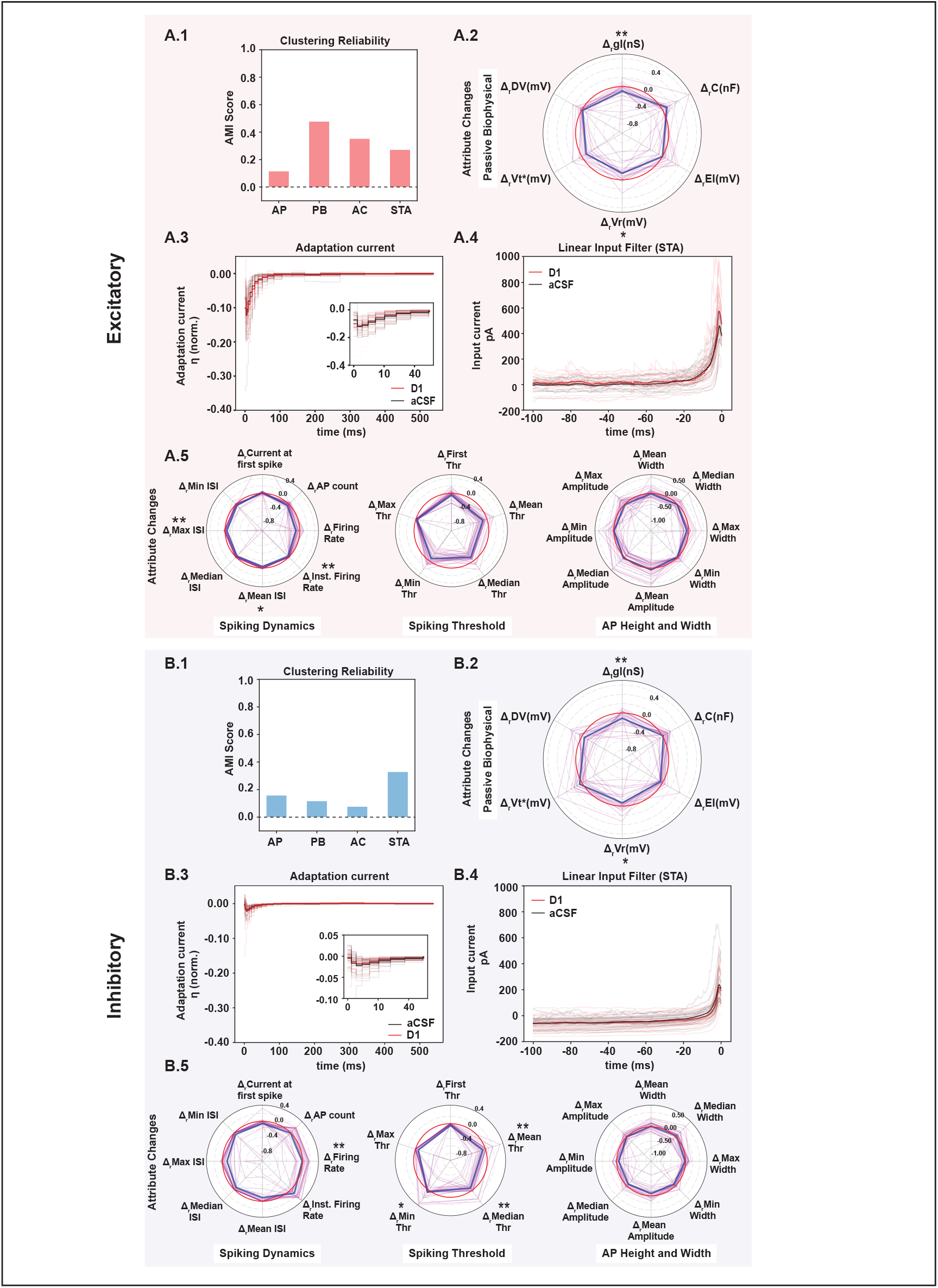
D1-R activation changes functional clustering for both excitatory and inhibitory neurons. (**A.1**) The adjusted mutual information score between the clustering labels obtained for control and D1 trials. (**A.2**) Change in passive parameters with respect to control as a result of D1 receptor activation, normalized by the control trial values. (**A.3**) Adaptation current for control (black) and D1 (red), the mean is represented with darker curves. (**A.4**) The spike triggered average profile for control (black) and D1 (red) trials, the mean is represented with a thick line. (**A.5**) Change in action potential attribute sets with respect to control as a result of D1 receptor activation, normalized by the control trial values. With the mean marked represented with a thick line and the zero line is coloured in red. (**B.1-5**) Same as **A.1-5** but for inhibitory neurons. * *p* 0.05, ** *p* ≤ 0.01, *** *p* ≤ 0.001.

Having established a stable baseline, we proceeded to cluster the neurons based on features explained above for each neuromodulator condition. For each agonist (D1, D2, M1), we applied dimensionality reduction (UMAP) and community detection (Louvain clustering) to the four attribute sets during control and agonist conditions separately (see Methods). We then measured the similarity in neuron classification before and after neuromodulation using the adjusted mutual information (AMI) between the control and agonist cluster labels. An AMI score near 1 indicates that clustering is unchanged by neuromodulation, whereas a low score indicates that neurons have been re-grouped into different functional clusters during neuromodulation condition. Next, we applied this framework to each agonist condition to quantify changes in functional clustering.

### D1 receptor activation drastically alters functional clustering

During D1-R agonist condition, the clustering of neurons based on each attribute set was markedly different from the control clustering. In excitatory neurons, the histogram of AMI scores for D1 vs. control (Figure 2A.1) was skewed towards low values for all four attribute sets (AP, PB, AC, STA). This indicates that D1-R activation caused many excitatory cells to switch their functional grouping, neurons that were similar during control conditions became dissimilar during D1-R activation, and vice versa, across multiple aspects of physiology. A similar outcome was observed for inhibitory neurons (Figure 2B.1), with consistently low AMI scores during D1-R activation for AP, PB, AC, and STA features. Thus, D1 modulation fundamentally reshuffled the functional identity of cells in both excitatory and inhibitory populations. In contrast, performing the same clustering comparison between two control trials yielded a much higher similarity than D1-R agonist condition (Figure S3), emphasizing that the shifts seen with D1 are due to receptor activation rather than random variability.

All three neuromodulators induced significant reorganization of clustering, but the extent and pattern were receptor-specific. D2-R and M1-R activation also produced low cluster similarity scores (AMI) between control and drug conditions, indicating that these modulators, like D1, redefine the functional groupings of neurons (Figure S2). However, D1 had the largest sample size and showed the clearest effects, so we focus on D1 as an illustrative case in the main text (full results for D2 and M1 are provided in Figure S4 & S5). Overall, across excitatory and inhibitory neurons, neuromodulator application consistently disrupted the original clustering structure based on intrinsic and evoked features, suggesting that receptor-specific modulation can create new functional subtypes or states that were not present for baseline conditions.

What neuronal properties drive these clustering changes? To gain insight, we examined how each functional attribute set was altered by neuromodulation, starting with the D1 receptor (which had the most substantial effect). We compared the distributions of individual features during control vs. D1 agonist conditions for both excitatory and inhibitory groups. For excitatory neurons during D1-R activation, we found notable changes in passive membrane properties (PB). In particular, input conductance (*g*_*L*_) and membrane reset potential (*V*_*r*_) were both significantly reduced by D1-R activation (one-sided t-tests; *p <* 0.001 for *g*_*L*_, *p <* 0.05 for *V*_*r*_; Figure 2A.2). A lower conductance and a more depolarized reset point generally indicate increased intrinsic excitability (cells become more responsive to current input). Surprisingly, other properties of excitatory neurons showed relatively little change with D1-R activation. Neither the adaptation current (AC) magnitude nor its kinetics were significantly affected by D1-R activation in excitatory cells: the time constants and peak amplitude of the adaptation currents were similar between control and D1 conditions (no significant differences in AC rise or decay; Figure S9). Likewise, the spike-triggered average (STA), which reflects the linear input filtering, remained essentially unchanged in excitatory neurons with D1-R activation: the STA peak and time-to-peak showed no significant differences between control and D1 (Figure S10). These results suggest that, for excitatory neurons, D1 modulation primarily impacts passive excitability properties, while leaving the shape of the adaptation current and the preferred input features (STA) intact.

In inhibitory neurons, D1-R activation altered passive properties and had a minimal effect on adaptation currents and STA, but with a few differences in detail. Similar to excitatory cells, inhibitory neurons showed a significant reduction in membrane conductance and a small hyperpolarizing shift in reset potential (*p <* 0.001 and *p <* 0.05, respectively, in Figure 2B.2) due to D1-R activation. We observed that adaptation currents and STA filters in inhibitory neurons were largely unchanged during D1-R activation (Figure 2B.3, 2B.4), the time constants and peak values of inhibitory adaptation currents showed no significant differences between control and D1-R conditions (paired t-tests, *p >* 0.4 for both decay and peak). However, the STA of inhibitory cells did show a minor change in timing: the STA rise time became slightly faster with D1 (*p≈* 0.02 for rise time), although the STA peak amplitude did not significantly change (*p≈* 0.08; inhibitory STA results in Figure S10). In summary, D1 receptor activation tended to increase intrinsic excitability (via passive properties) without dramatically altering the adaptive currents or preferred input features in both inhibitory and excitatory neurons.

We wanted to understand if there are differences between subgroups of neurons in the way their action potential and passive biophysical properties are altered due to D1-R activation. For this, we clustered the changes in AP and PB attributes between control and D1 trials for both excitatory and inhibitory neurons and summarized our findings using polar plots with each set of attributes in Figure 3. Each curve represents a single neuron, colored for its respective cluster identity and the mean is represented with a thick line. There are 3 clusters of AP differences for excitatory neurons and 4 clusters for inhibitory neurons. Similarly, we find 3 clusters each for PB differences for both excitatory and inhibitory neurons. We compared cluster assignments derived from control trials with those obtained by clustering the agonist-induced changes (agonist–control) using adjusted mutual information (AMI), to test whether neurons that group together for control conditions also exhibit similar responses to neuromodulation shown in the histogram at the bottom of Figure 3. AMI scores for excitatory and inhibitory neurons were consistently low across all 4 attributes, suggesting that changes in neuronal properties due to D1-R receptor activation were not organized according to the cell types defined by clustering of neurons based on the control trials. We performed a similar analysis for D2 and M1 agonist trials and summarized our findings in Figure S12 and S13.

**Fig. 3.**
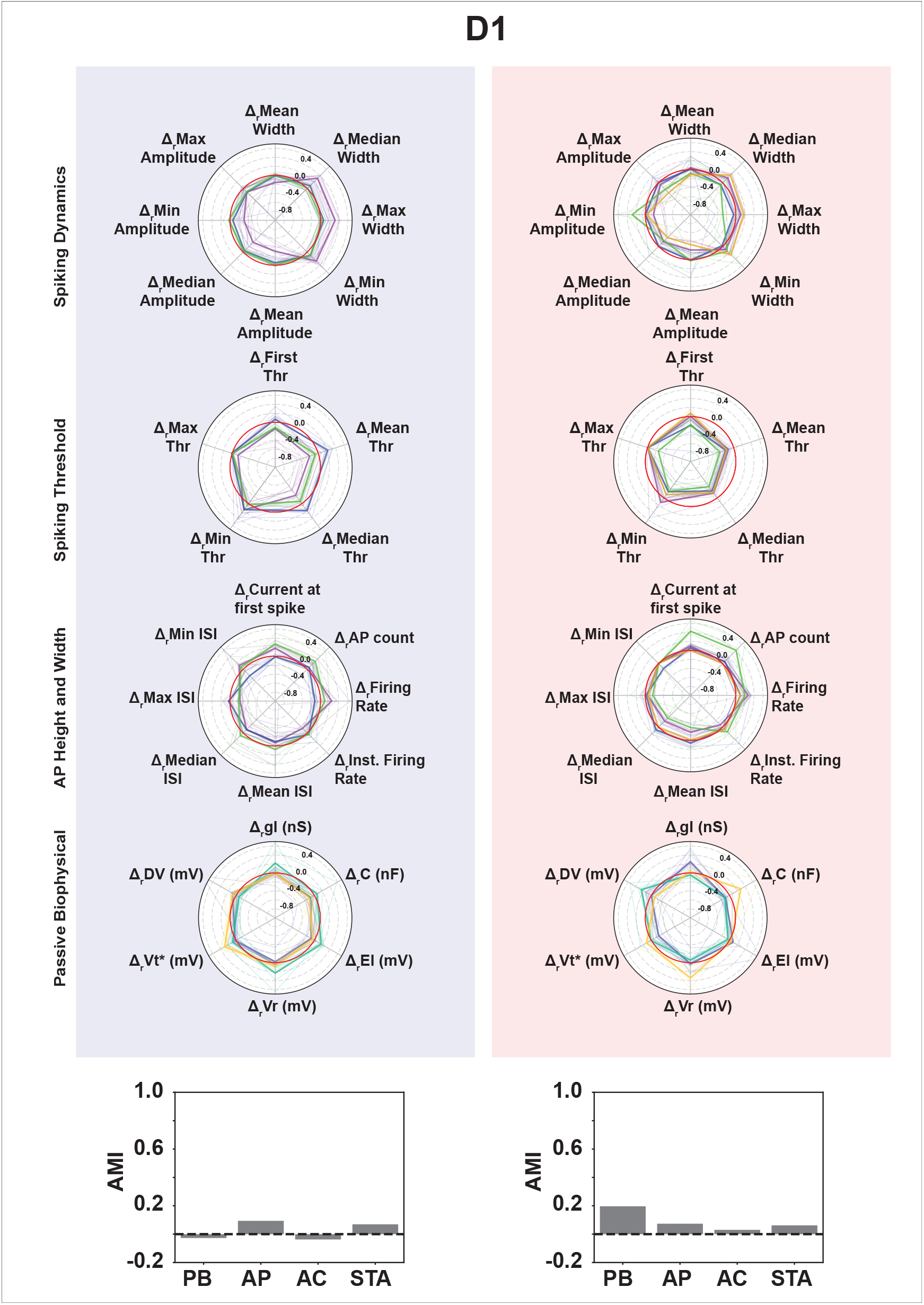
Clustering based on differences between control and D1 trials for action potential and passive biophysical attributes reveals subgroups of neurons getting modulated differently as a result of D1-R activation. Clusters based on difference values between D1 and control trials for action potential (subdivided into spiking dynamics, spiking threshold and AP height and width) and passive biophysical properties for both, excitatory (red background) and inhibitory neurons (blue background). Each neuron is represented with a thin line and colored with their respective cluster label. The mean for each cluster is represented with a thick line. The histogram at the bottom shows the AMI score between cluster results using control trials and the cluster results based on the difference between D1 and control trials.

The major effect of D1, then, was a reconfiguration of how combinations of attributes define cell identities, rather than significant changes in each attribute independently. This was further supported by analyzing the relationships among attributes during D1-R activation. We performed cosine similarity analyses on the shapes of adaptation current (AC) and STA curves, comparing within-condition similarity to across-condition similarity. For both, the AC and STA curves, we found that the similarity of responses was significantly higher when comparing within the same condition (two D1 trials or two control trials) than when comparing across conditions (control vs D1) (Kruskal–Wallis tests, *p <* 0.001; with post hoc tests showing within-condition *>* across-condition similarity; see Figure S11). In other words, STA and AC curves were more consistent within a given neuromodulatory state than between states. This consistent shift across the population (rather than random drift) underscores that D1 modulation reliably pushes neurons toward new response profiles. Notably, the within-D1 similarity was slightly lower than within-control for adaptation currents, implying that D1 modulation might also introduce some increased heterogeneity in AC responses across neurons, but in general both features showed a coherent population-wide change during D1-R activation. The net conclusion is that D1 receptor activation causes a systematic remodeling of neuronal functional characteristics across the board, reinforcing the idea that neuromodulation creates a distinct “network state” at the single-cell level.

Similar analyses for D2 and M1 receptor activation (Figure S11) revealed that each neuromodulator induces a unique pattern of attribute changes and clustering effects. In brief, D2-R activation had a qualitatively similar effect to D1 in many respects but generally weaker: for example, D2-R activation reduced excitatory membrane conductance, but the shifts were less pronounced than D1-R activation case. M1-R activation produced a distinct combination of effects, as we will describe in the context of covariance structure below. Importantly, all three receptor manipulations significantly altered functional clustering (Figure S2), indicating that receptor-specific neuromodulation consistently reshapes how neurons are functionally grouped, even though the detailed changes in attributes differ within subgroups of each cell-type.

### Neuromodulation reorganizes correlations among neuronal functional attributes

The clustering results above suggest that neuromodulators do not act on each neuronal property in isolation; instead, they alter the coordination between properties. To investigate this more rigorously, we employed a Multi-set Correlation and Factor Analysis (MCFA) (22) to quantify how the covariance structure between the four functional attribute sets changed for each neuromodulator condition. MCFA decomposes the variance in each attribute set into three components: shared variance (the variance in that set that is correlated with variance in other sets), private variance (the variance unique to that set, uncorrelated with others), and residual noise variance (22). By comparing these components between control and drug conditions for each cell type, we can determine whether neuromodulation increases or decreases feature coupling (i.e., shared variance) or independence (i.e., private variance). Since the sample size between drug and control trials were different, we apply a bootstrapping method to account for the imbalance (see Methods). Standard deviations (or confidence intervals) are included in Tables S6 & S6 to reflect the variability in control estimates.

Figure 4 illustrates the MCFA results as stacked variance components for each attribute set, in control vs. agonist conditions. We found that neuromodulation indeed produced receptor- and cell-type-specific restructuring of variance sharing. During D1-R receptor activation, for example, the proportion of private variance in excitatory neurons increased for the STA features (input selectivity), while the private variance for AP, PB, and AC attributes decreased relative to control (meaning those features became more shared). In other words, D1 made excitatory neurons’ input selectivity more independent from their other properties, but simultaneously tied their output-related properties (spiking dynamics and adaptation) more tightly together. In inhibitory neurons, D1 had the opposite broad effect: the private variance fractions for nearly all attributes decreased, and shared variance increased across the board (except perhaps a small exception for AC). This implies that D1-R activation drives inhibitory neuron properties to become more globally coordinated; their AP, PB, STA, and AC features all moved in a more lock-step fashion during modulation, resulting in a more integrated functional state. Quantitatively, the ratio of shared-to-private variance increased for inhibitory cells during D1-R activation for every attribute set, indicating a shift toward more cohesive, less independent features.

**Fig. 4.**
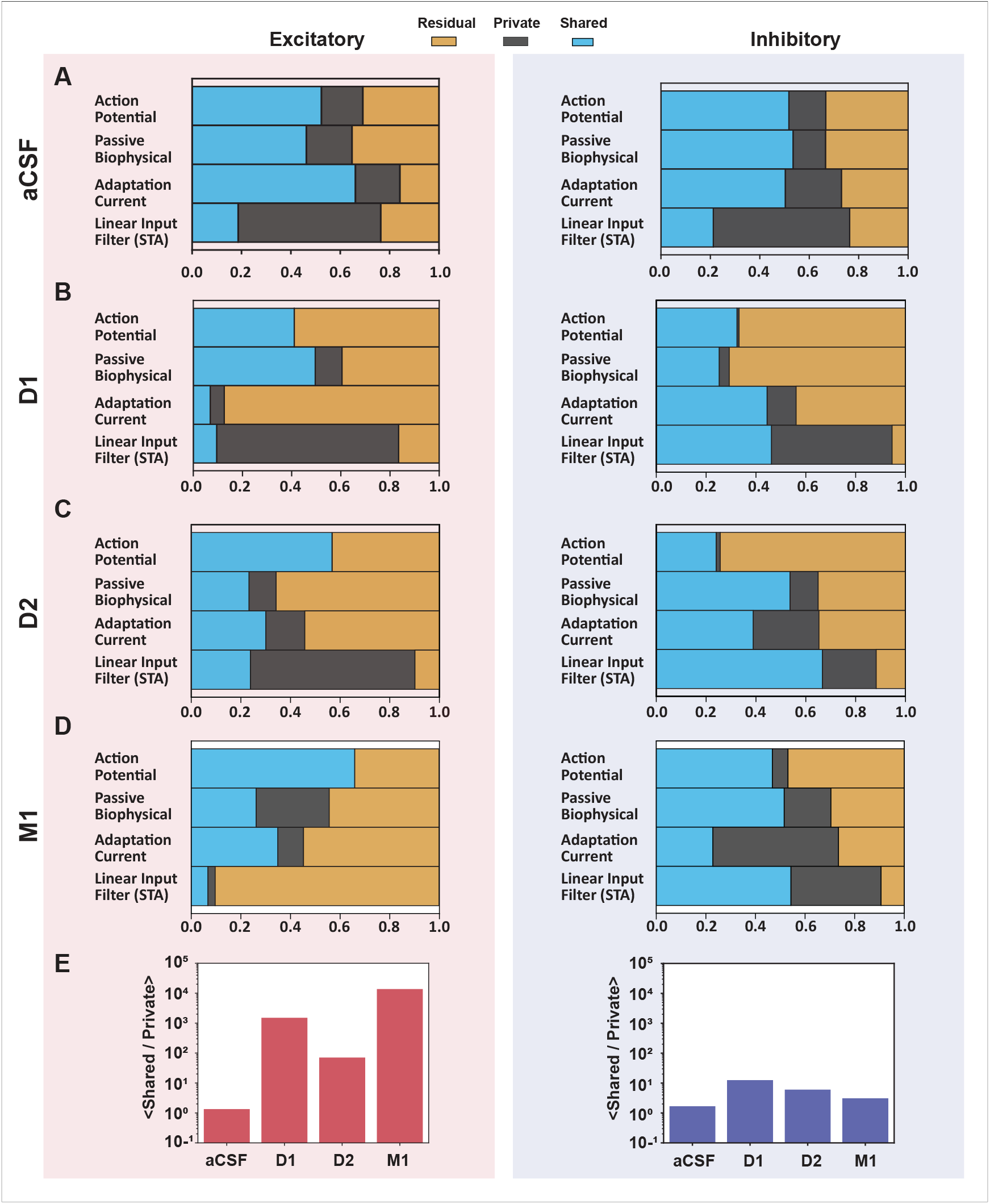
Correlation between functional attributes changes as a result of D1, D2 and M1 receptor activation compared to control in both excitatory and inhibitory neurons. **(A)** MCFA results for control-excitatory trials on the left and on the right is the control-inhibitory trials. **(B)** MCFA results of D1-excitatory trials on the left and the corresponding inhibitory trials is on the right. **(C-D)** Excitatory and inhibitory MCFA results for D2 and M1 agonist trials respectively. Excitatory trials are on the left and inhibitory on the right. **(E)** Histogram of the ratio of shared and private variance for control and each agonist conditions in the excitatory population. Similarly right histogram shows the ratio of shared over private variance for inhibitory population for each agonist condition.

The D2 dopamine receptor produced a similar qualitative pattern to D1 in our MCFA analysis, though more subtle. In excitatory neurons, D2 activation significantly raised the private variance of the STA set (even more than D1) and increased the shared variance between AP and AC features (Table S6). At the same time, D2, like D1, caused large drops in private variance for AP, AC, and PB attributes in excitatory cells (Table S6). The result was that excitatory the STAs became even more independent during D2-R activation than during D1-R activation, while AP and AC became more coupled (with each other and with other features). In inhibitory neurons, D2’s effect was straightforward: the private variance for all attributes (except AC) decreased and shared variance increased for all attributes (except a slight decrease for AP, see Table S7). In short, D2-R activation also drives inhibitory neuron properties to be more coupled (though perhaps not quite as uniformly as D1). The net effect of D2, much like D1, was cell-type specific: excitatory neurons showed selective decorrelation of STAs (indicating that input selectivity could change independently), whereas inhibitory neurons showed an across-the-board increase in coordination among features.

Muscarinic M1-R activation revealed yet another distinct pattern of reorganization. In excitatory neurons, M1 caused the shared variance to increase for AP and AC attributes (similar to D2’s effect on those features), while decreasing the shared variance for PB and STA. In other words, during M1-R agonist condition, excitatory spike output features (AP and adaptation) became more correlated with each other, but the passive membrane properties and input filtering became more independent. Additionally, M1 uniquely increased the residual (unexplained) variance for the STA in excitatory cells. We suspect this may be related to M1’s strong suppression of firing rate, which could introduce more “noise” in estimating the STA due to low number of spikes. For inhibitory neurons, M1’s effect was broadly similar to D1 and D2: shared variance increased for most attribute sets (AP, PB, STA) and private variance dropped for all except AC. Thus, M1-R activation pushed inhibitory neurons toward more coordinated properties (except that adaptation currents in inhibitory cells became somewhat more independent, a minor deviation).

In summary, across all receptors examined, neuromodulation tended to increase the global coordination of functional features in inhibitory neurons and to selectively re-balance the coordination in excitatory neurons. Dopamine D1 and D2 signaling in excitatory cells specifically decoupled the input selectivity (STA) from other traits, effectively allowing the kind of stimulus a neuron prefers to become a more independent degree of freedom, while making output excitability attributes (AP and AC) more interdependent. Cholinergic M1 modulation in excitatory cells also enhanced coupling among output-related properties (AP and AC) but promoted independence in passive and input-related properties. A comprehensive summary of these findings for each receptor and cell type is provided in Table S6 & S7 (summarizing changes in shared vs. private variance with detailed numerical values of variance components in cell-type specific manner). Table S8 & S9 and Figure S14 provide a summary of baseline MCFA analysis between control trials 1 and 2.

Crucially, the ratio of shared to private variance increased for every neuromodulator condition tested, for both excitatory and inhibitory populations, relative to the control condition. This indicates that neuromodulators generally act to enhance the coordinated variance in the system. Even when certain features (like STA in excitatory cells) become more independent, other features become more correlated, such that overall there is a shift toward more shared variability. Intuitively, one can think of this as neuromodulation reducing the dimensionality of variability in inhibitory neurons (making their properties move together, potentially reducing noise), while in excitatory neurons neuromodulation reassigns variance between features (promoting specialization of some features at the expense of others). These diverse strategies by which neuromodulators reshape the functional architecture of neurons suggest different computational roles: inhibitory neurons during neuromodulation may achieve more stable, coherent encoding (through integrated, tightly correlated properties), whereas excitatory neurons may gain more selective or flexible tuning (through the relative independence of certain features like input filtering).

## Discussion

In this study, we set out to investigate how neuromodulation, specifically activation of dopamine D1, dopamine D2, and muscarinic M1 receptors, alters the functional properties of cortical neurons beyond traditional measures of excitability. Using a frozen noise stimulus and whole-cell recordings from layer 2/3 putatively pyramidal (excitatory) cells and interneurons (inhibitory) in mouse somatosensory cortex, we quantified both the information transfer of single neurons and a broad set of intrinsic and synaptic response attributes during control and neuromodulator conditions. Our findings demonstrate that neuromodulators do far more than uniformly scaling up or down neuronal activity. Instead, they reconfigure the landscape of neuronal information processing in a cell-type and receptor-specific manner.

### Neuromodulators as selective regulators of encoding

We found that activating D1 and M1 receptors significantly reduced the fraction of sensory input information encoded by excitatory neurons, whereas it largely spared (or even slightly increased) the information encoded by inhibitory neurons. In practical terms, this is possibly indicative of a shift in the balance of coding: under the effect of certain neuromodulators, inhibitory neurons carry relatively more of the sensory information while excitatory neurons carry less. This result extends previous observations of D1 effects on firing rates (27–29) by showing that in the sensory cortex, D1 activation not only modulates spike rates but also diminishes the encoding efficiency of excitatory neurons. Interestingly, our finding that D1 and M1 dampen excitatory neuron encoding contrasts with a common expectation that neuromodulators broadly enhance excitability and thus might increase information throughput. Instead, the picture is more nuanced: dopamine and acetylcholine can redistribute information-processing roles among cell types. D2 receptor activation had a milder impact, slightly reducing excitatory encoding and moderately affecting firing, which might reflect its known role in subtly modulating excitability (9). Overall, the differential effects on excitatory vs. inhibitory neurons align with the idea that neuromodulators can target specific cell populations within cortical circuits to fine-tune network function (27, 30–32).

### Reshaping of functional identity

A key discovery of this work is that neuromodulation reshapes how neurons are functionally classified. During baseline conditions, neurons can be grouped into functional classes based on intrinsic electrophysiological features, consistent with prior taxonomy studies (33, 34). Our unsupervised clustering confirmed that, for example, neurons with similar spike shapes and membrane properties form broad clusters that correspond to known cell types (e.g., fast-spiking putatively inhibitory vs. regular-spiking putatively excitatory cells). However, when neuromodulators were introduced, sub-groups within these broad classes were reassigned across clusters indicating a reorganization of functional similarity rather than a loss of canonical cell-type structure. Neurons that were functionally similar before diverged into different clusters, and vice versa, implying that the neuromodulatory state can redefine a neuron’s identity (different from cell-type) in functional terms. This finding is reminiscent of context-dependent classification schemes (21, 25, 26), and here we show that the context of neuromodulation is enough to recast neuronal identity. Our clustering analysis revealed that all three receptor-specific neuromodulations (D1, D2, M1) disrupted the original clustering of neurons based on multiple feature sets. This suggests that the effect is quite general: any significant neuromodulatory input can create a new functional configuration of the circuit. Notably, inhibitory neuron clusters were as affected as excitatory ones, meaning interneurons also shift their functional relationships during modulation. Overall, our clustering results suggest that neuromodulators do more than gain modulation to neurons; they dynamically restructure the computational topology of cortical circuits, thereby reassigning neurons to new functional roles as neuromodulatory tone changes.

### Coordination vs. independence of neuronal properties

By using MCFA, we gained insight into how these functional reassignments occur. Neuromodulators altered the correlation structure among neuronal attributes, rather than uniformly affecting each feature independently. A striking pattern emerged: inhibitory neurons consistently moved toward a state of greater coordination across properties, whereas excitatory neurons gained independence in specific properties (e.g., input selectivity) during D1-R and D2-R activation. From a computational perspective, increasing shared variance among inhibitory neuron properties could lead to more constrained and coherent firing patterns, effectively reducing internal noise and variability. This makes sense if we consider inhibitory interneurons as providing a stable, synchronized regulatory influence on the network; neuromodulators may enhance that role by aligning their functional parameters. Indeed, our data show an increased shared variance (and reduced private variance) in inhibitory cells for all neuromodulators, suggesting that neuromodulation could make inhibitory processing more robust and stereotyped across the population. Conversely, in excitatory neurons, neuromodulation (especially via D1 and D2) made certain features more independent, in particular, it uncoupled the neurons’ stimulus selectivity (STA) from their other intrinsic properties. This increase in private variance for STA means that during neuromodulation, an excitatory neuron’s preferred input features can vary more freely, without being tightly linked to its other traits. We speculate that this could enable greater diversity in feature tuning in excitatory cells, potentially enriching the dimensionality of sensory representations. In other words, dopamine might enable excitatory neurons to explore more varied stimulus-response mappings by loosening the constraints that normally tie their input filtering to their overall excitability profile. Such flexibility could be advantageous for adaptive coding in changing contexts, albeit at the cost of reduced immediate efficiency (as evidenced by the drop in information transfer for excitatory cells). Muscarinic modulation presented a somewhat different case. M1 activation increased coupling of some features (AP & AC) for excitatory neurons while freeing others (PB, STA). This mixed strategy suggests that acetylcholine may bias excitatory neurons toward more feature-selective representations, strengthening certain response aspects while decorrelating others (28, 29). Meanwhile, M1 made inhibitory cells more coordinated overall. It is tempting to relate this to known effects of acetylcholine on cortical networks, such as enhancing signal-to-noise ratio and sharpening tuning in sensory processing (2, 3). Our results provide a mechanistic view: by increasing shared variance (coordination) in inhibitory neurons, acetylcholine could dampen uncorrelated noise, and by modulating coordination in excitatory neurons, it could refine which stimulus features are represented. Building on the MCFA results, we propose three network-level predictions that can be tested directly in vivo. First, the increased low-dimensional coordination observed within inhibitory populations predicts that neuromodulatory receptor activation will stabilize inhibitory gain control, yielding reduced trial-to-trial variability in inhibitory output and more consistent regulation of population excitability during repeated sensory drive. Second, the relative independence of excitatory stimulus selectivity (STA) from other intrinsic domains predicts that excitatory ensembles will show greater flexibility in retuning when task demands or contextual variables shift, enabling rapid remapping of stimulus representations without requiring commensurate changes in broader intrinsic excitability or spike dynamics. Third, because neuromodulators reconfigure shared and private variance across functional domains, combined receptor engagement (e.g., D1 and M1 co-activation) should produce non-additive interactions, including synergy or occlusion, manifesting as disproportionately large (or unexpectedly constrained) changes in population coding, coupling structure, and information transfer relative to either receptor manipulation alone.

### Implications for cortical computation and disease

Collectively, our results suggest that neuromodulators act not only as simple gain controllers but as high-level regulators of neuronal computation. By increasing the proportion of shared variance among neuronal properties in inhibitory cells, neuromodulators may reduce internal noise and enforce more uniform, reliable processing, supporting robust and coherent network activity. This could underlie cognitive states requiring focused, stable processing (for instance, dopamine’s role in working memory may involve stabilizing inhibitory microcircuits). In contrast, by increasing private variance in certain excitatory features, neuromodulators might promote high-dimensional, flexible representations in excitatory networks, which could be important for exploratory or learning states (allowing excitatory neurons to explore new encoding schemes). Thus, neuromodulation can bias the cortex toward either more integrated, noise-resistant coding (inhibitory-driven) or more specialized, diverse coding (excitatory-driven), depending on the context. These findings are relevant to understanding disorders involving neuromodulatory dysfunction. For example, in schizophrenia and Parkinson’s disease, dysregulation of dopamine is central to the development of the clinical phenotype (10–12). Our results imply that an imbalance in D1/D2 receptor activity could lead to misalignment between excitatory and inhibitory processing, perhaps causing excitatory neurons to carry too little information or become mis-tuned, and inhibitory neurons either over-constraining the circuit or failing to stabilize it. Additionally, the receptor-specific differences we observed (D1 vs. D2 vs. M1) underscore that therapies targeting neuromodulators must consider the nuanced effects on circuit computation, not just overall firing rates.

### Limitations and future directions

The present work examined the neuromodulation of single neurons in an in-vitro setting. While this allowed precise control and measurement of intrinsic properties, it provides only a partial view of how neuromodulators affect circuit dynamics in vivo (where network feedback and spatiotemporal neuromodulator release patterns play a role). Future work should extend these findings to intact circuits and behavioral contexts to confirm that these single-cell changes translate to network-level computational changes. Second, we focused on single receptors in isolation. Neuromodulatory systems often act in concert, and it will be important to investigate how multiple neuromodulator signals interact (e.g., co-activation of D1 and M1) to shape neuronal function. Our analysis provides a baseline for single-receptor effects, which could be built upon to study combined modulation. Third, our information measurements were based on a specific frozen noise stimulus and assumed stationarity (ergodicity) over a few minutes of recording. It is possible that different input statistics or longer timescales of neuromodulator action could reveal additional effects. It is also worth noting that two types of FN inputs were used during the recordings based on the responsiveness of the neurons (see (21) Methods), which could have had an effect on our findings. Nonetheless, the consistency of our results across many cells and the clear systematic changes give confidence in the main conclusions.

In conclusion, we have shown that cell-type specific neuromodulation fundamentally reshapes neuronal information transfer and the coordination among neuronal properties. Rather than acting uniformly, neuromodulators impart a rich, structured reorganization, stabilizing inhibitory neuron function and diversifying excitatory neuron function. This orchestrated modulation likely underlies neuromodulatory systems’ ability to support flexible cognition, allowing cortical circuits to reconfigure on the fly in response to varying computational demands.

## Methods

A detailed explanation of data collection and analysis methods is provided in the supplementary materials.

### Ethics statement

The data used in this research is made freely available to the community. All experimental work, as described in the articles cited, was carried out in accordance with the European directive 2010/63/EU, the Dutch national regulations and international standards for animal care and use.

### Slice electrophysiology

The experimental procedures were described before (20, 35) and in the supplementary materials. In short, slices were prepared from adult mice expressing Cre recombinase under the control of either the parvalbumin promoter (RRID: MGI:5315557) or the somatostatin promoter (RRID: IMSR JAX:013044), backcrossed to the C57BL/6 background. The animals were housed in a 12-hour light / dark cycle with ad libitum access to food and water and kept in family cages until the day of the experiment.

The experimental procedures followed previously published protocols (36–40). Coronal slices (thickness: 300 microns) were cut from the barrel cortex subregion of the primary somatosensory cortex, and individual neurons were visualized using differential interference contrast (DIC) optics at 40× magnification before whole-cell access using pipettes with a resistance of 5-9 *M* Ω. The internal (pipette) solution contained (in mM): 130 K-gluconate, 5 KCl, 1.5 *MgCl*_2_·6*H*_2_*O*, 0.4 *Na*_3_*GTP*, 4 *Na*_2_*ATP*, 10 HEPES, 10 Na-phosphocreatine, and 0.6 EGTA. The pH was adjusted to 7.22 with KOH. Recordings were performed using HEKA EPC10 amplifiers and PatchMaster software (v2×90.2).

Targeted pharmacological activation of the select receptors was performed using muscarinic acetylcholine M1 receptors (McN-A-343, 20*·µM*), dopamine D1 (SKF38393, 1*·µM*) and D2 receptors (Quinpirole, 10*·µM*). All compounds were purchased from Sigma-Aldrich and dissolved in ACSF (41).

## Supporting information

Methods and supplemental items

## Data Availability

All recorded current clamp data and the code to analyze them can be found in this repository: https://doi.org/10.34973/nhes-qe97. The database is an extended version of the dataset (35, 42), specifically with neuromodulation. The code and extracted feature files for all figures and analyses is available here: https://github.com/Nishant-codex/single_cell_analysis.git.

## ACKNOWLEDGEMENTS

This project has received funding from the European Union’s Horizon 2020 research and innovation program under the Marie Skłodowska-Curie grant agreement No 860949.

