## Supplementary material for "Neuromodulatory Control of Cortical Function: Cell-Type Specific Regulation of Neuronal Information Transfer": Methods and supplemental items

DRAFT

### This PDF contains:

Methods Details
Supplementary Figures S1-S14
Supplementary Tables S1-S8
SI References

DRAFT

**Materials and Methods.** First, we quantified the per-cell change in FI and FR between drug and control trials:

$$\Delta_r \mathrm{FI}_{\mathrm{ago}} = \frac{\mathrm{FI}_{\mathrm{agonist}} - \mathrm{FI}_{\mathrm{aCSF}}}{\mathrm{FI}_{\mathrm{aCSF}}}, \quad \Delta_r \mathrm{FR}_{\mathrm{ago}} = \frac{\mathrm{FR}_{\mathrm{agonist}} - \mathrm{FR}_{\mathrm{aCSF}}}{\mathrm{FR}_{\mathrm{aCSF}}}$$

We compared these values between excitatory and inhibitory neurons to assess whether receptor-specific neuromodulation affects cell types differently. To account for intrinsic trial-to-trial variability, we also computed $\Delta_r$FI and $\Delta_r$FR across two control (aCSF) trials from the same recording sets.

$$\Delta_r \mathrm{FI}_{\mathrm{aCSF}} = \frac{\mathrm{FI}_{\mathrm{aCSF}_2} - \mathrm{FI}_{\mathrm{aCSF}_1}}{\mathrm{FI}_{\mathrm{aCSF}_1}}, \quad \Delta_r \mathrm{FR}_{\mathrm{aCSF}} = \frac{\mathrm{FR}_{\mathrm{aCSF}_2} - \mathrm{FR}_{\mathrm{aCSF}_1}}{\mathrm{FR}_{\mathrm{aCSF}_1}}$$

**Frozen Noise (FN) protocol** The Frozen Noise input protocol consisted of injecting a somatic current that is the result of an artificial neural network of 1000 neurons responding (firing Poisson spikes) to random stimuli i.e., the hidden state, the membrane potential response to the somatic input is recorded with a sampling rate of 20 kHz for a total length of 360 seconds and saved. Each raw data file consisted of a vehicle control trial (artificial Cerebrospinal fluid i.e. aCSF) and a drug trial (a specific neuromodulatory receptor agonist or antagonist was added to the bath and the recording was repeated). Some files consisted of multiple control and drug trials. See (1, 2) for more details. A schema for the protocol is shown in Fig. 1. In total 288 neurons (Table. 1) were analyzed, we discarded recording sets with high levels of noise.

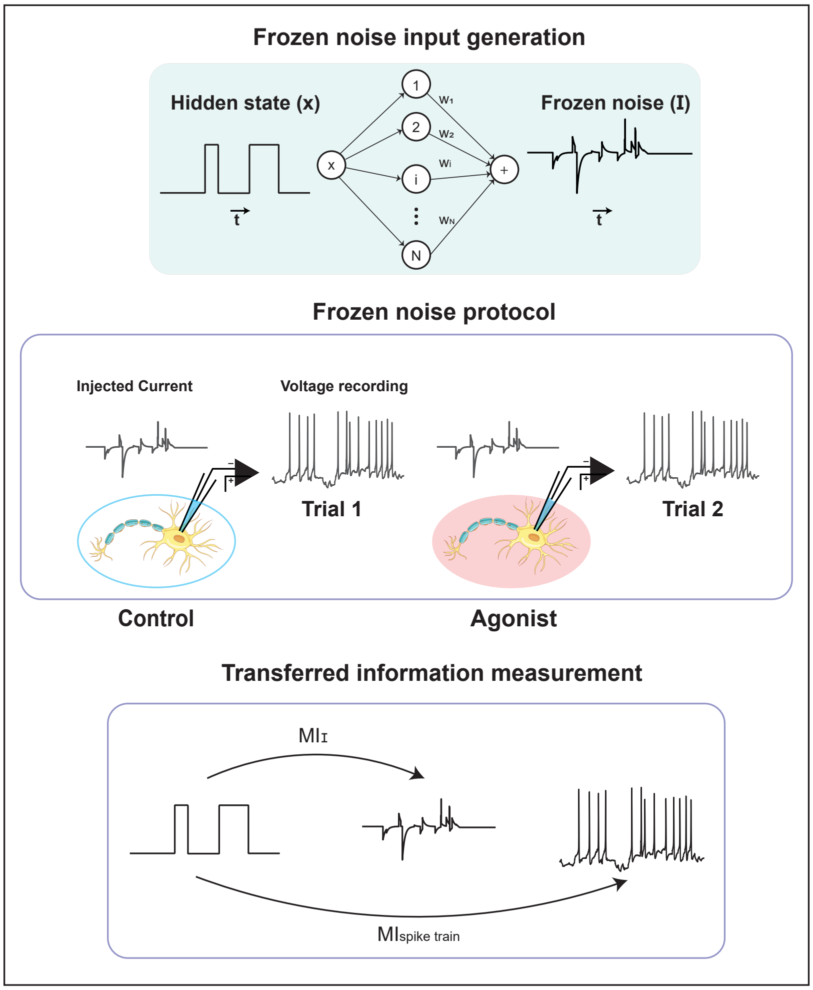


**Fig. 1. Frozen noise protocol**: The top figure provides an illustration of how input is generated, the middle figure illustrates how this input is injected into the soma of a neuron for both aCSF and agonist conditions, the bottom figure illustrates the schema for calculating mutual information.

| Condition | Cell-type | Trials |
|---|---|---|
| aCSF | Excitatory | 183 |
| aCSF | Inhibitory | 121 |
| D1 | Excitatory | 50 |
| D1 | Inhibitory | 36 |
| D2 | Excitatory | 40 |
| D2 | Inhibitory | 19 |
| M1 | Excitatory | 19 |
| M1 | Inhibitory | 20 |

**Table 1.** Number of recording set for control and agonist conditions

### Analysis.

#### *Feature Extraction.*

We extracted waveform, action potential, passive biophysical and spike triggered average from single neuron recordings recorded under Frozen Noise input protocol, these recordings were performed first under a vehicle control (aCSF) and then repeated with a receptor agonist added to the bath.

**Spike waveforms** As explained in (3), we identified peaks from the membrane potential traces and kept the hyperparameters and ISI threshold criteria the same as in (3). The length of the waveforms used in this study is 10ms (5ms before and after the peak).

**Action potential attributes** The action potential attributes were extracted to study the dynamics, threshold and waveforms related attributes throughout a trial via descriptive statistics. The action potential attributes were extracted for aCSF and agonist trails as described in (3). A summary of all the Action potential attributs is provided in Table. 2.

***Passive Biophysical Feature extraction using GLIF model.*** In order to extract passive biophysical attributes as well as adaptation current from aCSF and agonist trials, we fit a Generalized Leaky Integrate and Fire (GLIF) neuron model (4) as described in detail in (3). We take the first 100 second from the recording as the training set to extract passive attributes as well as adaptation current from the recording. A summary of all the passive biophysical features extracted is provded in Table. 3. Using the same GLIF model fitted to neural recordings, we also extracted the adaptation current $\eta(t)$ triggered by a spike event, see (3, 4).

**Spike Triggered Average** The spike-triggered average (STA) is the average shape of the stimulus that precedes each spike. We extracted the STA using the following equation given by (5):

$$STA = \frac{1}{N}\Sigma_{n=1}^{N}\vec{s}(t_n), \qquad \textbf{(1)}$$

where $t_n$ is the $n^{th}$ spike time, s is the stimulus vector preceding the spike for a fixed time window of 100 ms, and N is the total number of spikes. Before clustering, we standardize (i.e. z score) and then normalize the STA vector with an $L_2$ norm. We didn't use any kind of whitening or regularization to calculate the STA.

**UMAP + Louvain clustering.** Universal Manifold Approximator (UMAP) is a non-linear dimensionality reduction algorithm which is advantageous for preserving global structure of the data (6) in lower dimensions, this makes it more suitable for visualization especially for high dimensional datasets ($p >> N$, where p is the dimensionality of the data and N is number of samples) compared to other methods such as PCA which fail to perform due to curse of dimensionality (7). As explained in (3, 8) the high-dimensional graph obtained during the intermediate step in the UMAP algorithm can be exploited to perform clustering using Louvain community detection (9). We chose the hyperparameter based on cluster stability criteria and the corresponding number of clusters based on the same heuristic as explained in (3).
For measuring clustering similarity between aCSF and drug conditions, we calculate a cluster similarity matrix as explained in (3). For quantifying the similarity between labels assigned to aCSF versus drug trials, we used cluster similarity measure such as adjusted mutual information score using scikit-learn Python package (10). The AMI score is 1 when the two cluster labels are identical. Random labels can have an expected AMI around 0 on average and therefore can be negative.

**A. Information transfer protocol.** We employed the information transfer protocol detailed in (1, 11) to quantify how much information a single neuron extracts from its inputs. This protocol assumes that neurons respond to the presence or absence of a preferred stimulus, which is modeled as a binary hidden state ($\mathbf{x}$) that switches between 0 and 1 according to a memoryless Markov process. Importantly, the neuron does not directly observe this hidden state; instead, it receives input from a large

DRAFT

| Feature Group | Feature | Description | Summary Statistics |
|---|---|---|---|
| Spiking Dynamics | Current at first spike | Current amplitude at which the neuron first crosses threshold to fire a spike. | Single value per trial. |
| | AP count | Total number of action potentials generated during a trial. | Single value per trial. |
| | Time to first spike | Time (in ms) from trial start to first spike. | Single value per trial. |
| | Firing rate | Total number of spikes divided by trial duration (spikes/sec). | Single value per trial. |
| | Interspike Interval (ISI) | Time between successive spikes $(t_{n+1} - t_n)$. | Mean, median, max, min calculated. |
| | Instantaneous rate | Reciprocal of ISI $(1/(t_{n+1} - t_n))$. | Mean, median, max, min calculated. |
| Spike Threshold | Spike threshold | Voltage at which $dV/dt$ first exceeds 25 mV/ms. Calculated per spike. | Mean, median, max, min computed. |
| AP Height & Width | Width | Time from threshold crossing to return below threshold after AP peak. | Mean, median, max, min calculated. |
| | Amplitude | Voltage difference between AP peak and threshold. | Mean, median, max, min calculated. |

**Table 2.** Summary of Action Potential Attributes and Their Descriptions

DRAFT

| Feature | Description |
|---|---|
| Membrane Capacitance ($C$) | The cell membrane's ability to store charge; influences how quickly voltage changes in response to current. |
| Leak Conductance ($g_L$) | Governs the passive flow of ions across the membrane; contributes to the rate of membrane potential decay. |
| Resting Potential ($E_L$) | The membrane voltage the cell settles at in the absence of input; baseline membrane potential. |
| Sharpness of Spike Threshold ($\Delta_r V$) | Controls how sharply the firing probability increases as membrane potential approaches threshold. |
| Threshold Baseline ($V_T^*$) | The baseline value of the dynamic spike threshold; determines the average voltage required to trigger a spike. |
| Reset Voltage ($V_{reset}$) | The membrane potential value the neuron resets to after a spike and refractory period. |

**Table 3.** Passive Biophysical Features Extracted from the GLIF Model

population of simulated presynaptic neurons. Each presynaptic neuron fires a Poisson spike train, with firing rates $q^i_{on}$ when the hidden state is ON ($x = 1$), and $q^i_{off}$ when it is OFF ($x = 0$).
To generate the input current ($\mathbf{I}$), the spike train of each presynaptic neuron is convolved with a 5 ms exponential kernel and then weighted by $w_i = \log\left(\frac{q^i_{on}}{q^i_{off}}\right)$. The weighted signals are summed to produce the total input current, which is then scaled and injected as somatic input during in vitro patch-clamp recordings. The neuron's membrane potential and spike times are recorded in response to this current (Fig. 1).
Information transfer is quantified in several steps. First, the entropy of the hidden state ($H_{xx}$) is calculated. Next, the mutual information between the hidden state and the input current ($MI_I$), and between the hidden state and the neuron's spike times ($MI_{\text{spike times}}$), are computed. The fraction of information (FI) transferred by the neuron is then defined as:

$$FI = \frac{MI_{\text{spike times}}}{MI_I} \tag{2}$$

Since the entropy of the hidden state is always greater than or equal to the mutual information, $FI$ values range between 0 and 1. This approach assumes ergodicity (so time averages equal ensemble averages) and that spike trains are approximately Poissonian, though minor deviations from Poisson statistics do not significantly affect the results. By using a binary hidden state, this protocol allows for reliable estimation of mutual information even from relatively short recordings, providing a robust measure of how effectively a neuron's spikes encode information about dynamic stimuli.

**Quantification and statistical analysis.** We used the non-parametric Wilcoxon test to test the significance of the observed difference in information transfer (FI) between paired aCSF and drug trials . We also used one sided Student's T-test to test the significance of the observed change between aCSF and agonist condition for passive biophysical and action potential attributes. Significance value was set to $p < 0.05$ in both cases. We performed a Kruskal-Wallis H test and a post-hoc Mann-Whitney U test with Bonferroni correction for multiple comparisons for comparing the cosine similarity between aCSF and agonist trials. All statistical tests were performed using scipy-stats package (12).

**Multi-set Correlation and Factor Analysis.** To investigate how neuromodulation alters the relationship between different functional attribute sets (AP, PB, AC, and STA), we used **Multi-set Correlation and Factor Analysis (MCFA)** (13), an unsupervised integration method designed to model both shared and private variance across multiple high-dimensional data types from the same samples. This method combines elements of canonical correlation analysis and factor analysis to produce low-dimensional representations of common and dataset-specific structure.
Each attribute set was z-scored (mean-centered and variance-scaled), and MCFA was applied separately to aCSF and agonist conditions. For the aCSF condition, we fixed the number of principal components to 2 for all attribute sets, due to the relatively low dimensionality of AP and PB features (e.g., PB: 6 dimensions). Private latent dimensionality $k_m$ was also set to 1 for each set, based on model stability and convergence tests. Attempts to use higher latent dimensions resulted in non-convergent fits. These same dimensionality parameters (PCs = 2, $k_m$ = 1 ) were applied to agonist conditions to maintain consistency and due to smaller sample sizes in drug trials.
Since the number of aCSF recordings exceeded the number of agonist trials, we bootstrapped the control condition by randomly sampling subsets matched in size to the agonist group with the least number of recording sets, i.e., M1-R. This process was repeated 100 times, and MCFA was run on each subset. The resulting shared and private variances were averaged across runs to produce stable estimates for the control condition.
The MCFA fitting procedure used an expectation-maximization (EM) algorithm as described in (13), initialized according to (3). Convergence was assessed by monitoring the log-likelihood and change in loading matrices across iterations.

DRAFT

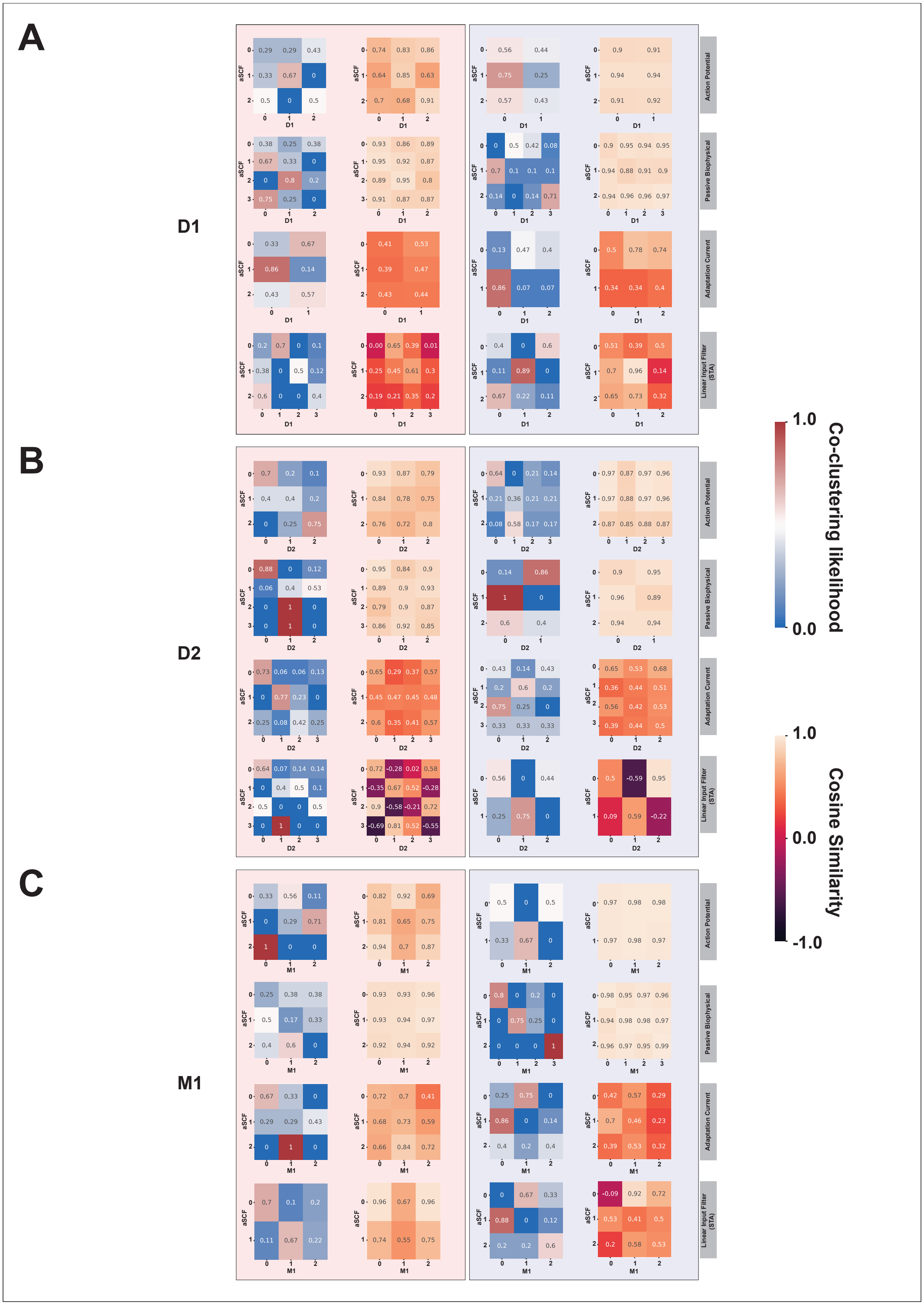


**Fig. 2. Clustering likelihood matrix and cosine similarity across labels shifts for all four properties as a result of neuromodulation for D1-R, D2-R and M1-R** Top panel shows the cluster likelihood and cosine similarity for D1 vs aCSF trials. Middle shows the cluster likelihood and cosine similarity for D2 vs aCSF trials. Bottom shows the cluster likelihood and cosine similarity for M1 vs aCSF trials.

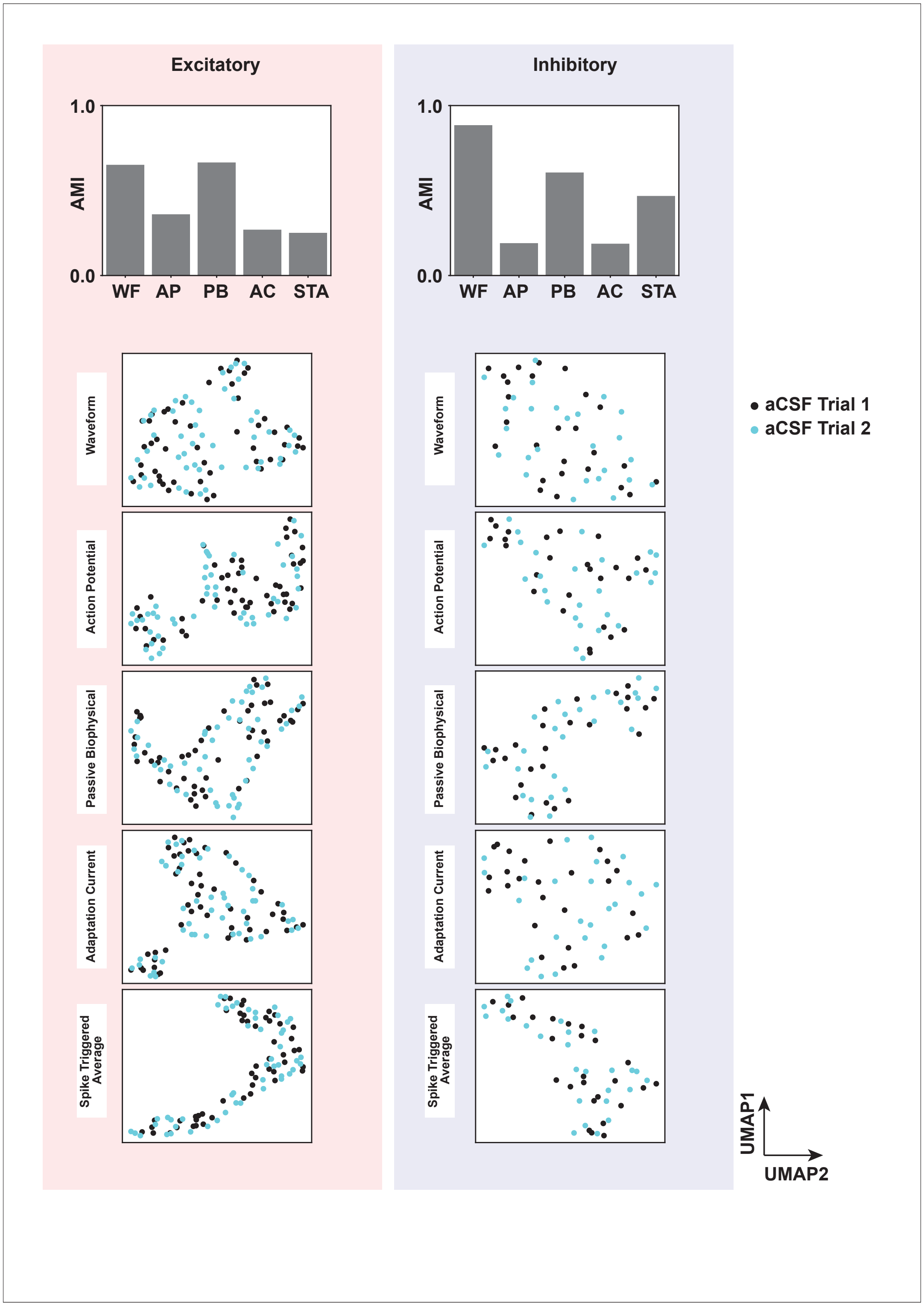


**Fig. 3. Comparison of aCSF trial 1 vs trial 2 clustering and manifold.** Left and right histograms on top the AMI score between cluster IDs found for aCSF trial 1 vs aCSF trial 2. Waveforms and passive biophysical properties were found to be consistent across the two trials as these properties are not strongly dependent input dynamics. The UMAP plots show the alignment for aCSF trial 1 vs aCSF trial 2.

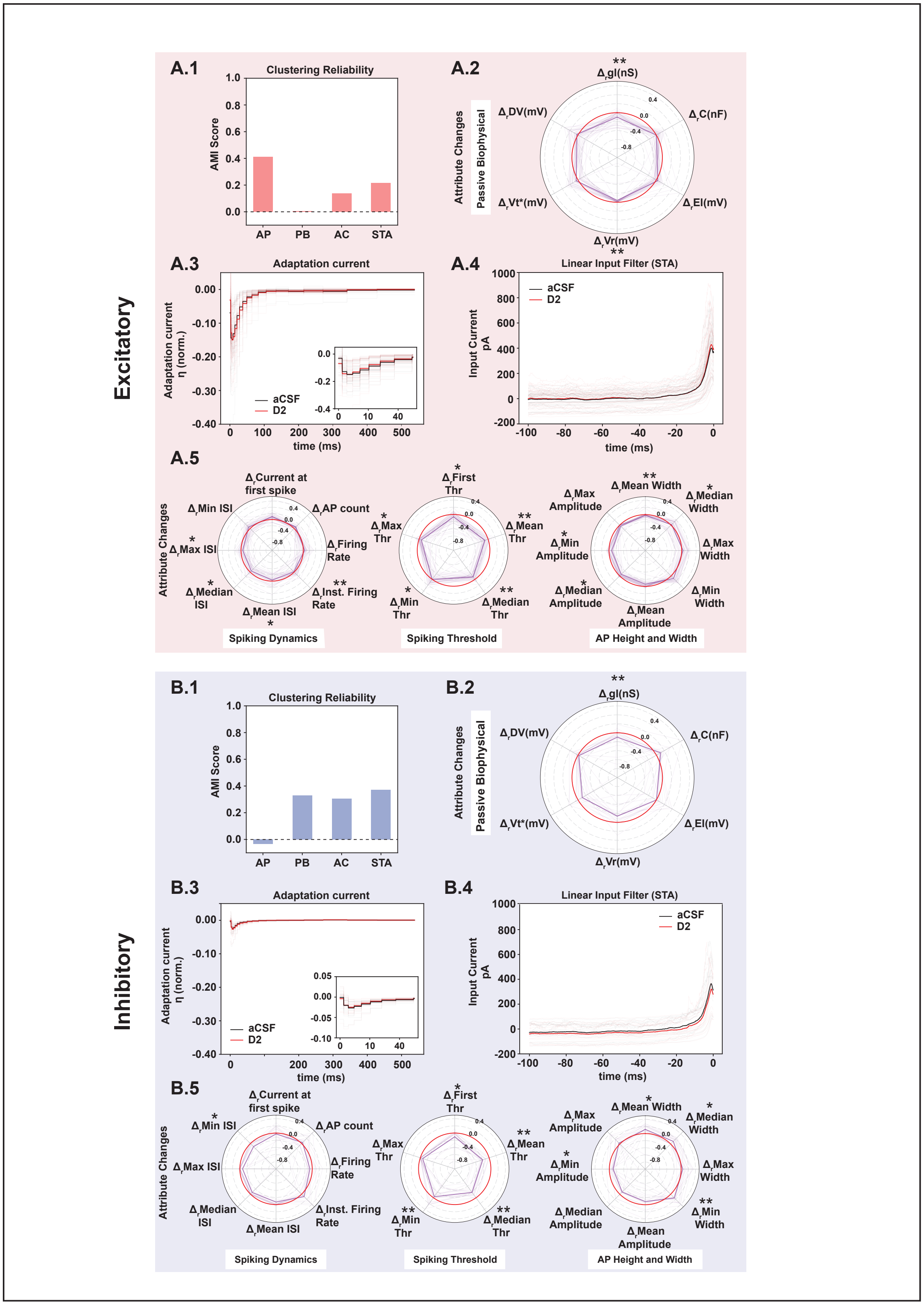


**Fig. 4. D2 receptor activation changes functional clustering and attributes for both excitatory and inhibitory neurons.** (**A.1**) Histogram shows the adjusted mutual information score between the clustering labels obtained for aCSF and D2 trials, the histogram shows a shift in cluster identities as a result of D2 receptor activation across four attributes. (**A.2**) shows the change in passive parameters with respect to control as a result of D2 receptor activation, normalized by the control trial values. With the mean marked represented with a thick line. (**A.3**) shows the adaptation current for control (black) and D2 (red), the mean is represented with darker curves. The adaptation current profiles between D2 and aCSF trials were found to be similar. The rise time as well the peak adaptation current between aCSF and D2 were found to be non-significant (see Appendix 9). (**A.4**) shows the spike triggered average profile for aCSF (black) and D2 (red) trials, the mean is represented with a thick line. (**A.5**) shows the change in action potential attribute sets with respect to control as a result of D2 receptor activation, normalized by the control trial values. With the mean marked represented with a thick line and the zero line is colored in red. The significant features are marked. **B.1-5** Same analysis repeated for inhibitory

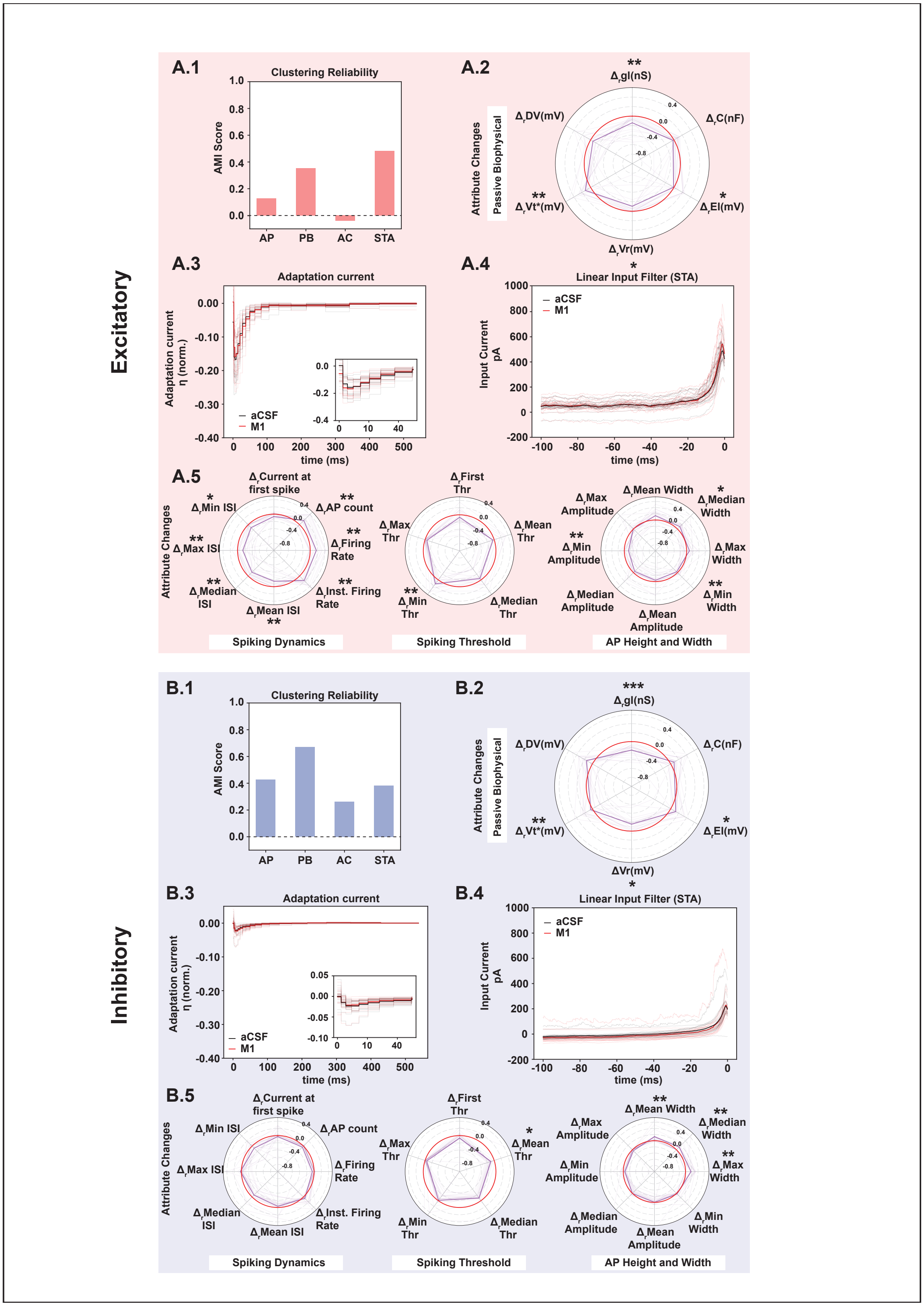


**Fig. 5. M1 receptor activation changes functional clustering and attributes for both excitatory and inhibitory neurons. A.1**) Histogram shows the adjusted mutual information score between the clustering labels obtained for aCSF and M1 trials, the histogram shows a shift in cluster identities as a result of M1 receptor activation across four attributes. (**A.2**) shows the change in passive parameters with respect to control as a result of M1 receptor activation, normalized by the control trial values. With the mean marked represented with a thick line. (**A.3**) shows the adaptation current for control (black) and M1 (red), the mean is represented with darker curves. The adaptation current profiles between M1 and aCSF trials were found to be similar. (**A.4**) shows the spike triggered average profile for aCSF (black) and M1 (red) trials, the mean is represented with a thick line. (**A.5**) shows the change in action potential attribute sets with respect to control as a result of M1 receptor activation, normalized by the control trial values. With the mean marked represented with a thick line and the zero line is colored in red. **B.1-5** Same analysis repeated for inhibitory neurons. * $p \le 0.05$, ** $p \le 0.01$, *** $p \le 0.001$.

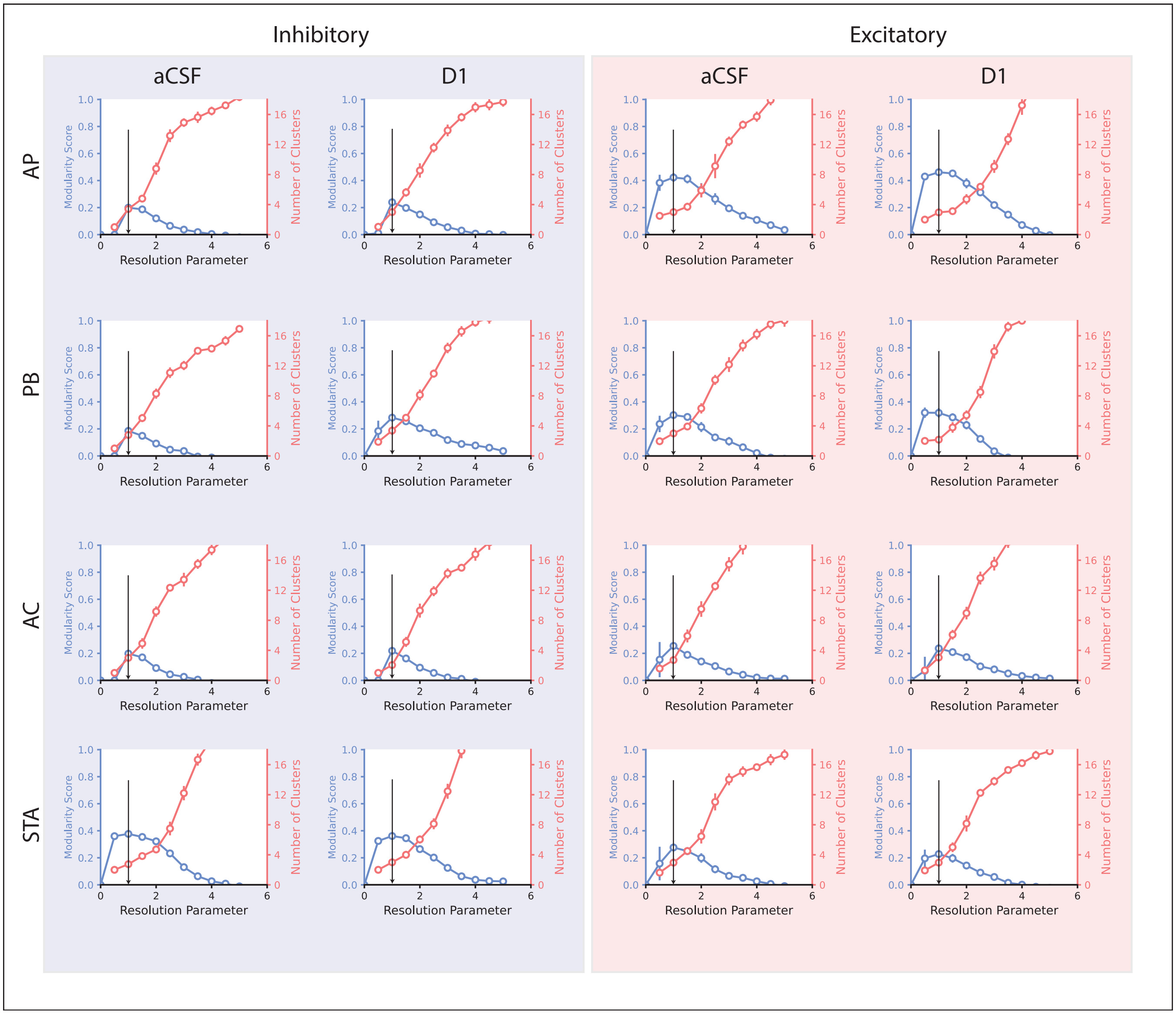


**Fig. 6. Stability of clusters over a range of resolution parameters and the corresponding number of clusters for D1** The plot shows the modularity score for each resolution parameter and the corresponding number of clusters. The selected resolution parameter is marked with the black arrow, this corresponds to the highest modularity score.

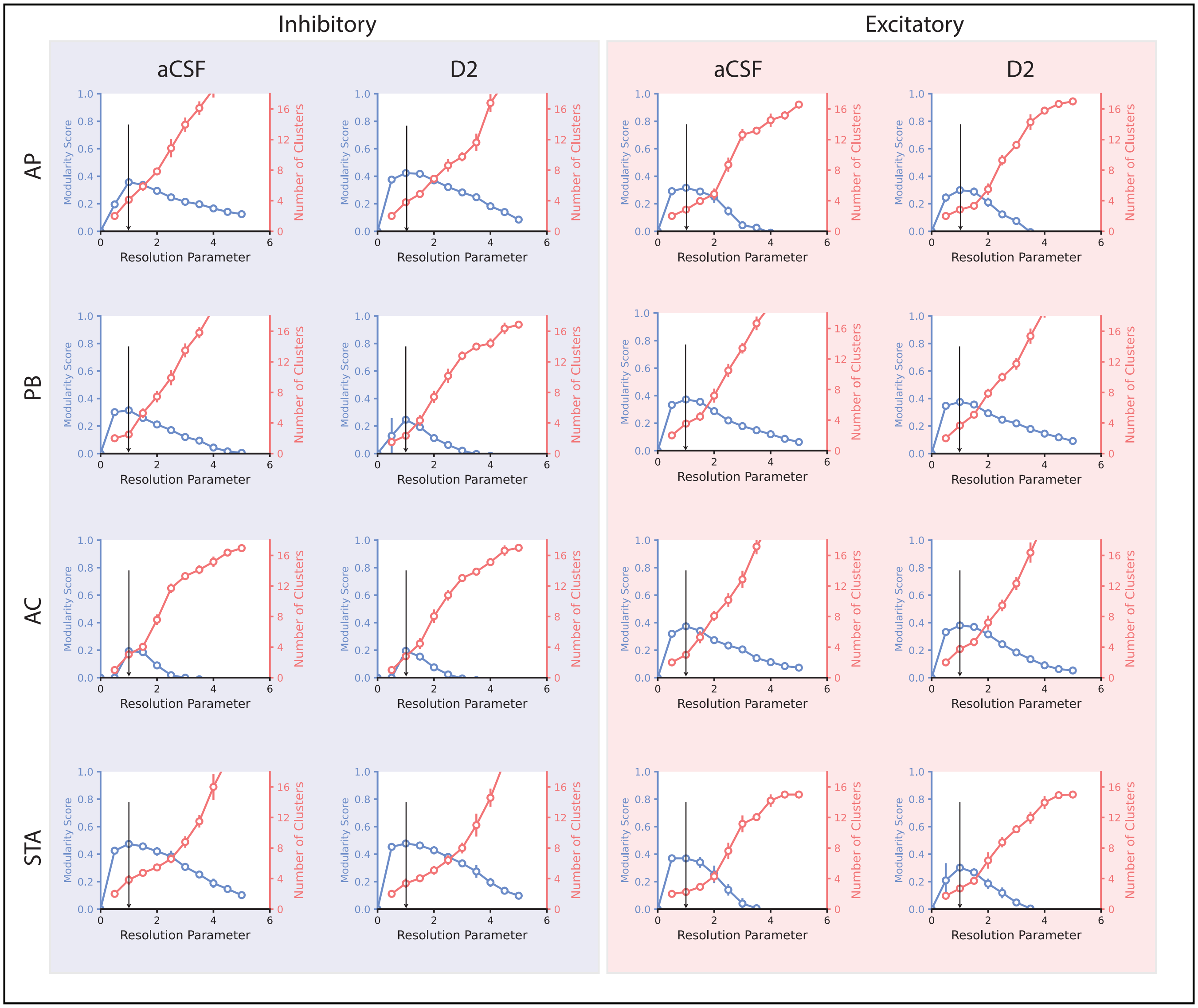


**Fig. 7. Stability of clusters over a range of resolution parameters and the corresponding number of clusters for D2** The plot shows the modularity score for each resolution parameter and the corresponding number of clusters. The selected resolution parameter is marked with the black arrow, this corresponds to the highest modularity score.

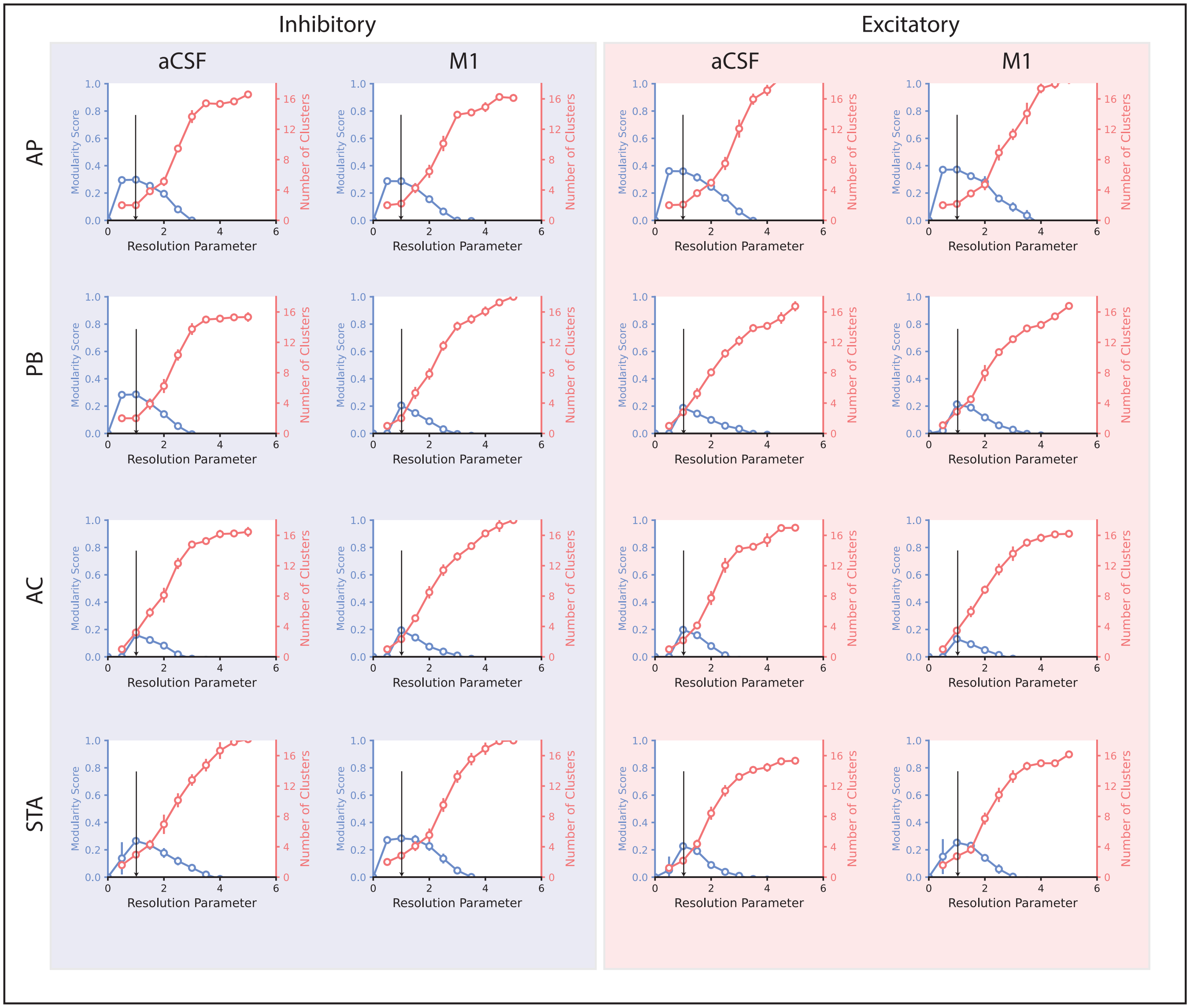


**Fig. 8. Stability of clusters over a range of resolution parameters and the corresponding number of clusters for M1** The plot shows the modularity score for each resolution parameter and the corresponding number of clusters. The selected resolution parameter is marked with the black arrow, this corresponds to the highest modularity score.

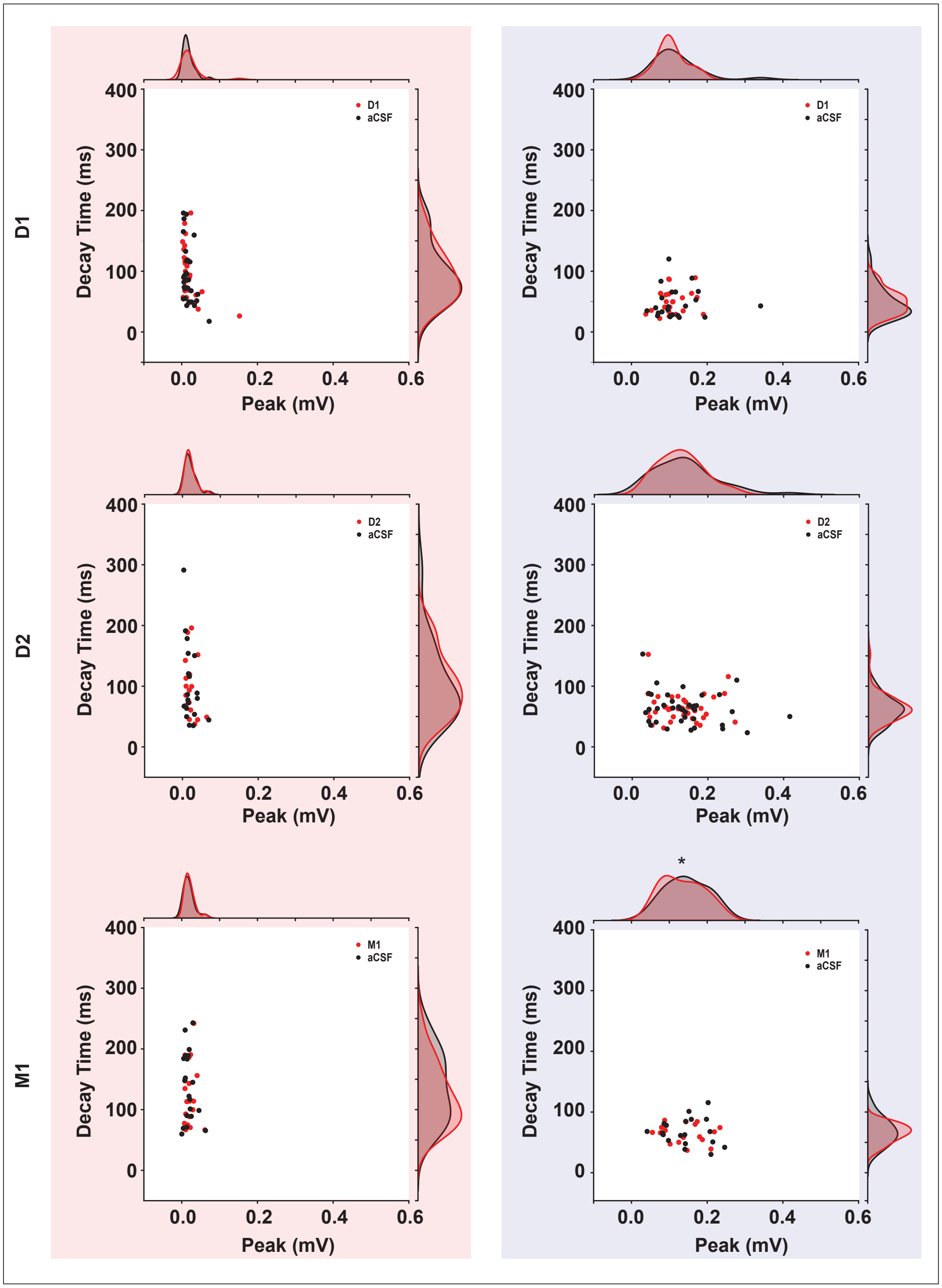


**Fig. 9. Adaptation current peaks and decay times are not strongly altered by D1, D2 and M1 modulation** The scatter plots show the normalized peaks of the AC curve on the x-axis and the decay time for D1, D2 and M1 vs aCSF trials. The red panel represents excitatory and blue panel represents inhibitory. * $p \leq 0.05$, ** $p \leq 0.01$, *** $p \leq 0.001$.

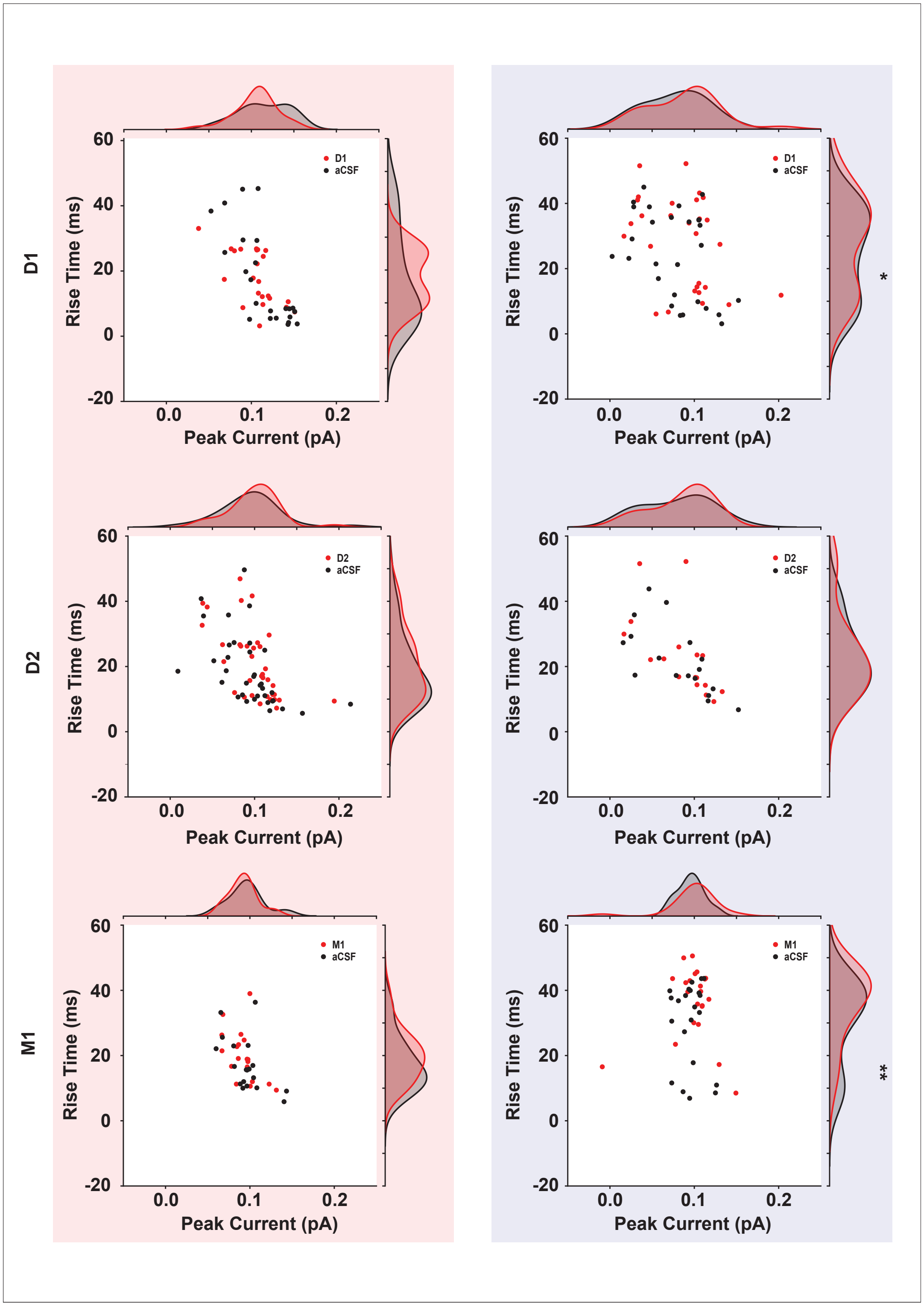


**Fig. 10. Spike triggered average peaks and rise times are modestly altered by D1, D2 and M1 modulation** The scatter plots show the normalized peaks of the STA curve on the x-axis and the rise time for D1, D2 and M1 vs aCSF trials. The red panel represents excitatory and blue panel represents inhibitory population. * $p \leq 0.05$, ** $p \leq 0.01$, *** $p \leq 0.001$

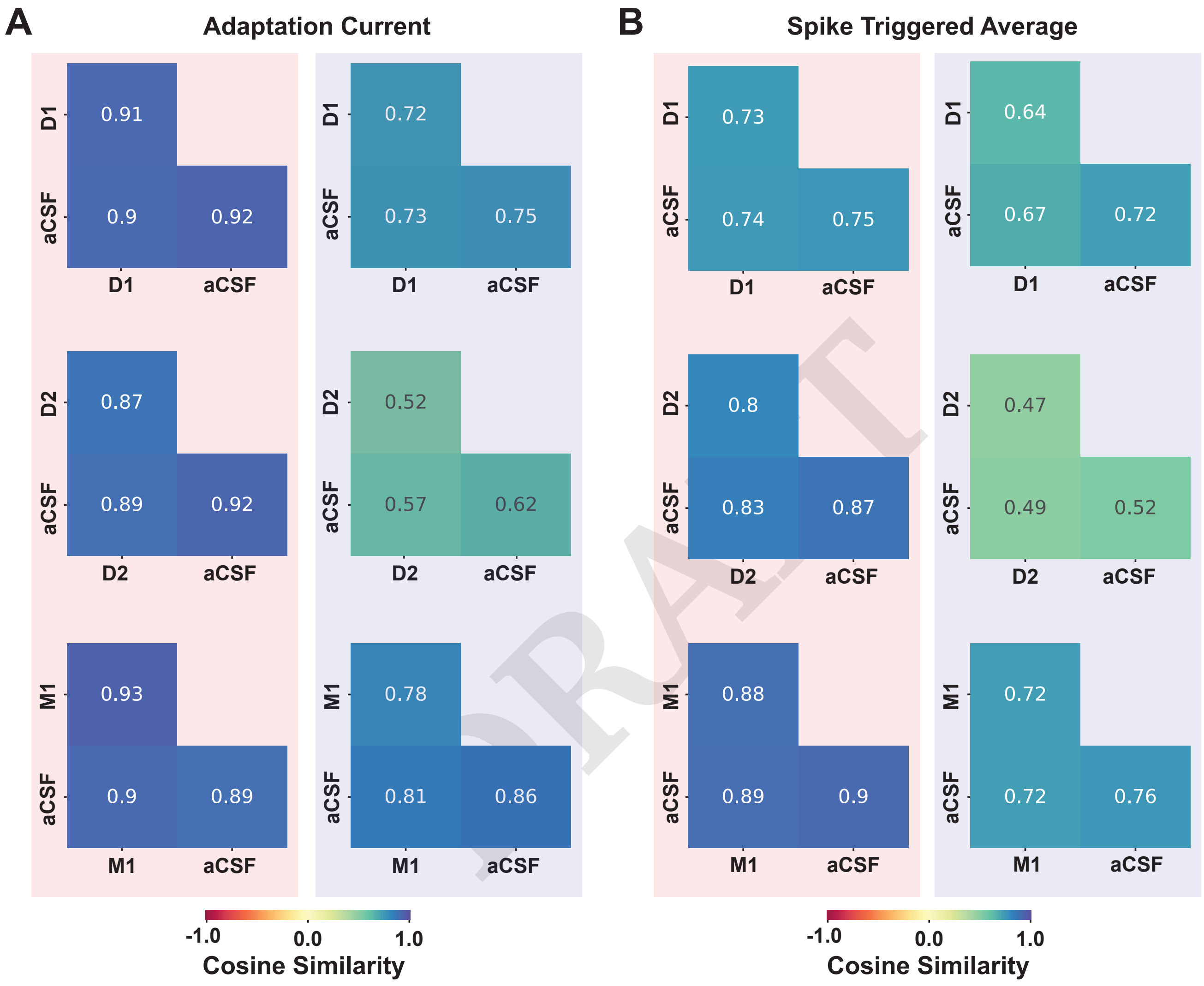


DRAFT

**Fig. 11. Average cosine similarity across and within aCSF-Agonist trials vary in a cell-type specific manner.** (**A**) Heatmaps show the average AC similarity for excitatory (red background) and inhibitory (blue background) for D1, D2 and M1 vs aCSF trials. (**B**) Heatmaps show the average STA similarities for excitatory (red background) and inhibitory (blue background) for D1, D2 and M1 vs aCSF trials.

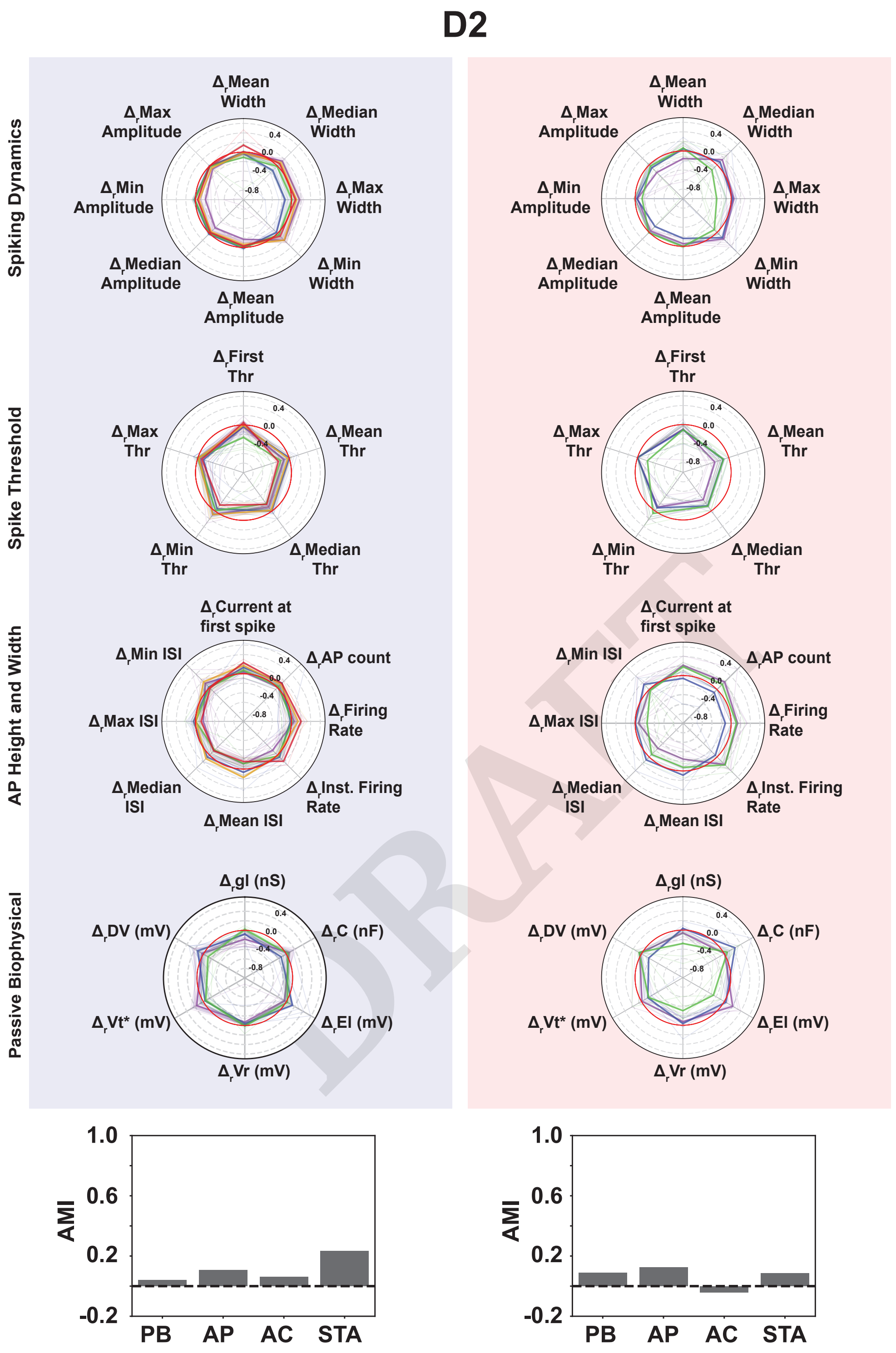


**Fig. 12. Clustering based on differences between control and D2 for action potential and passive biophysical attributes reveals subgroups of neurons getting modulated differently as a result of D2-R activation.** The polar plots show the clusters based on difference values between D2 and aCSF trials for action potential (subdivided into spiking dynamics, spiking threshold and AP height and width) and passive biophysical properties for both, excitatory (red background) and inhibitory (blue background). Each neuron is represented with a thin line and Coloured with their respective cluster label. The mean for each cluster is represented with a thick line. The histogram at the bottom shows the AMI score between cluster labels using aCSF trials and the cluster labels based on difference between D2 and aCSF trials.

DRAFT

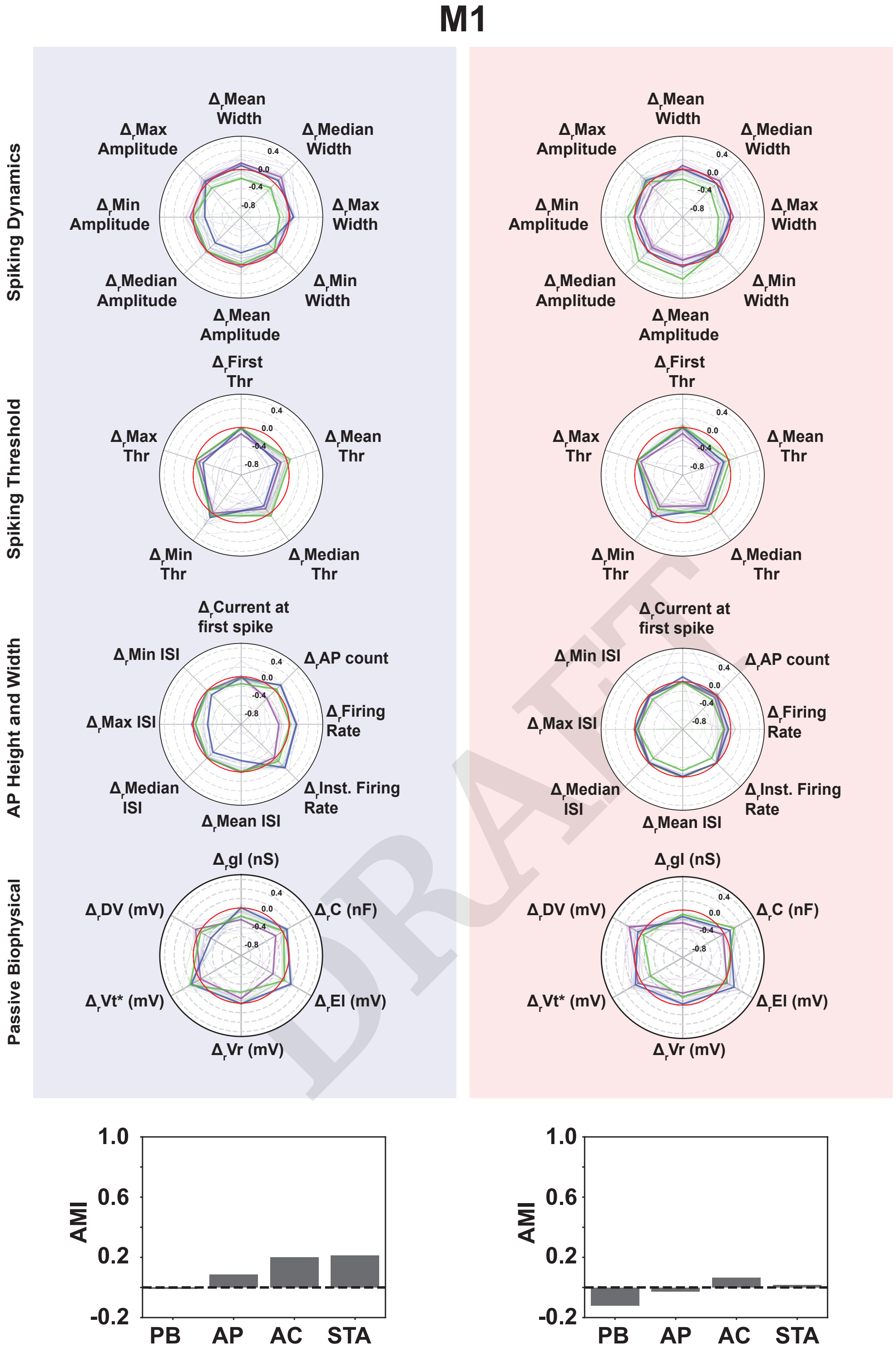


**Fig. 13. Clustering based on differences between control and M1 for action potential and passive biophysical attributes reveals subgroups of neurons getting modulated differently as a result of M1-R activation.** The polar plots show the clusters based on difference values between M1 and aCSF trials for action potential (subdivided into spiking dynamics, spiking threshold and AP height and width) and passive biophysical properties for both, excitatory (red background) and inhibitory (blue background). Each neuron is represented with a thin line and Coloured with their respective cluster label. The mean for each cluster is represented with a thick line. The histogram at the bottom shows the AMI score between cluster labels using aCSF trials and the cluster labels based on difference between M1 and aCSF trials.

DRAFT

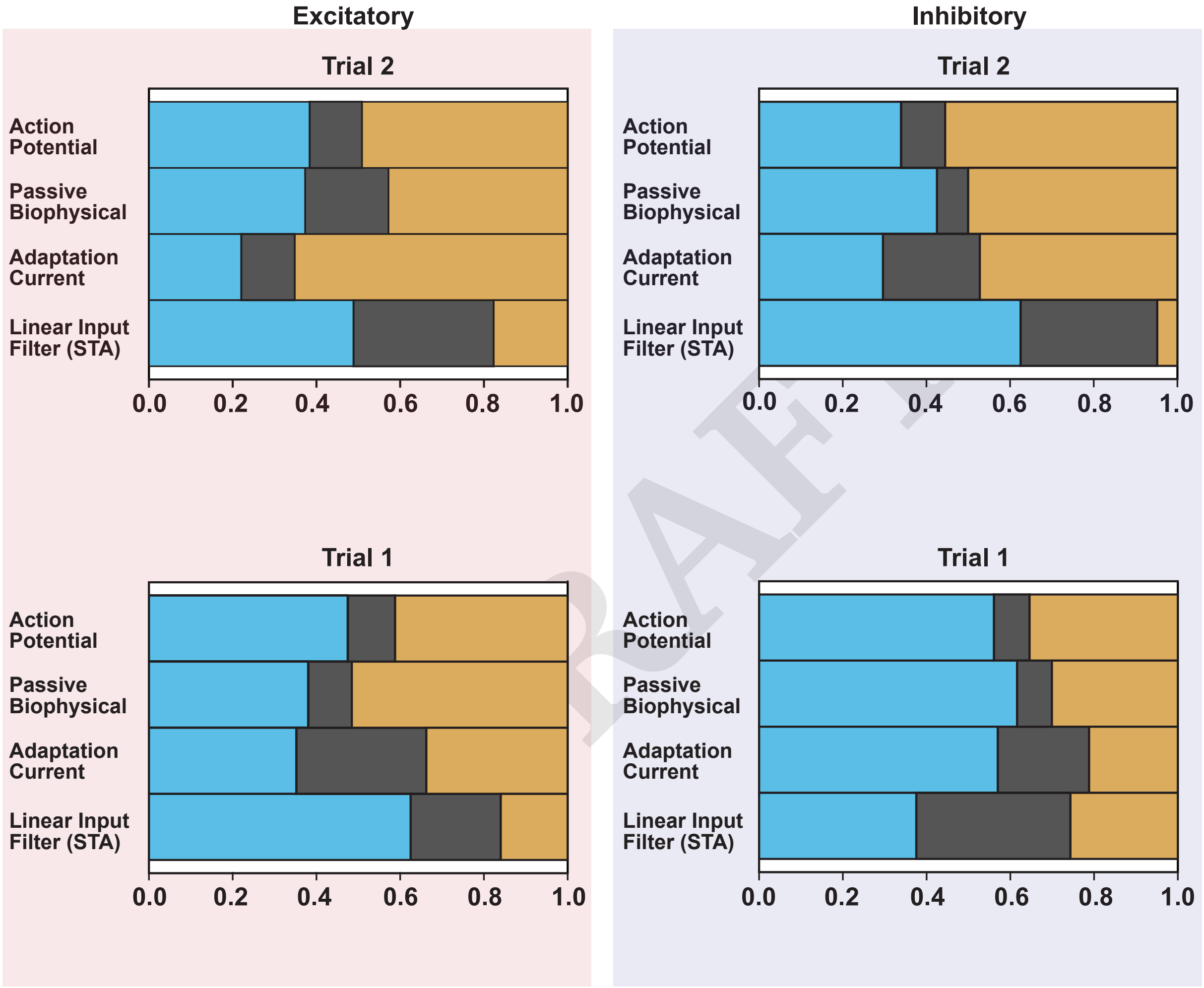


**Fig. 14. MCFA for 1st and 2nd aCSF trials.**

| Condition | Cell Type | Wilcoxon P-value | Statistic | Cohen's $d$ |
|---|---|---|---|---|
| D1–aCSF | Inhibitory | 0.019044 | 110 | –0.2409 |
| D1–aCSF | Excitatory | 0.104543 | 84 | 0.5385 |
| D2–aCSF | Inhibitory | 0.768005 | 87 | –0.0287 |
| D2–aCSF | Excitatory | 0.012016 | 225 | 0.3053 |
| M1–aCSF | Inhibitory | 0.523511 | 106 | –0.1181 |
| M1–aCSF | Excitatory | 0.002022 | 22 | 0.4790 |
| $aCSF_1$–$aCSF_2$ | Inhibitory | 0.205410 | 170 | –0.3161 |
| $aCSF_1$–$aCSF_2$ | Excitatory | 0.000111 | 340 | 0.3231 |

**Table 4.** Statistical comparison of fractional information (FI) changes under receptor activation.

DRAFT

| $\Delta FI$ | t-statistic | P-value | Cohen's-d |
|---|---|---|---|
| D1 | -3.16672 | 0.00270 | -0.94362 |
| D2 | -1.54012 | 0.12916 | -0.43694 |
| M1 | -2.95219 | 0.00538 | -0.95977 |
| $aCSF_2 - aCSF_1$ | -2.57383 | 0.01183 | -0.60172 |
| $\Delta FR$ | t-statistic | P-value | Cohen's-d |
| D1 | -3.66695 | 0.00062 | -1.09003 |
| D2 | -2.49095 | 0.01573 | -0.70850 |
| M1 | -3.20390 | 0.00274 | -1.04055 |
| $aCSF_2 - aCSF_1$ | -3.94022 | 0.00016 | -0.92093 |

**Table 5.** Statistical comparison of $\Delta FI$ and $\Delta FR$ between excitatory and inhibitory populations

DRAFT

| Attribute | Variance type | aCSF | D1 | D2 | M1 |
|---|---|---|---|---|---|
| AP | Private | 16.7742 (+/- 11.11) | 0.0062 | 0.1991 | 1.18E-03 |
| AP | Shared | 30.8543 (+/- 13.21) | 41.0578 | 55.0319 | 66.0431 |
| AP | Residual | 52.3714 (+/- 12.18) | 58.9360 | 44.7689 | 33.9556 |
| PB | Private | 18.5046 (+/- 13.50) | 10.4084 | 14.4461 | 29.4316 |
| PB | Shared | 35.2320 (+/- 15.97) | 49.5931 | 22.2190 | 26.3528 |
| PB | Residual | 46.2633 (+/- 15.84) | 39.9985 | 63.2628 | 44.2155 |
| AC | Private | 18.0218 (+/- 17.14) | 5.7216 | 16.8436 | 10.3262 |
| AC | Shared | 15.8369 (+/- 11.35) | 6.9056 | 30.6685 | 35.0802 |
| AC | Residual | 66.1412 (+/- 21.22) | 87.3728 | 44.7689 | 54.5935 |
| STA | Private | 57.7380 (+/- 24.75) | 73.9663 | 71.2200 | 2.8344 |
| STA | Shared | 23.6072 (+/- 19.86) | 9.5260 | 18.0047 | 6.9094 |
| STA | Residual | 18.6546 (+/- 18.49) | 16.5077 | 10.7753 | 90.2561 |

**Table 6. Excitatory MCFA results**. Values that are reduced compared to bootstrapped aCSF are colored in red and values that are increased compared to aCSF are colored in blue.

DRAFT

| Attribute | Variance type | aCSF | D1 | D2 | M1 |
|---|---|---|---|---|---|
| AP | Private | 14.9272 (+/- 09.5664) | 0.8297 | 1.6361 | 6.3213 |
| AP | Shared | 33.3099 (+/- 12.0733) | 32.3923 | 24.0248 | 46.7960 |
| AP | Residual | 51.7627 (+/- 10.2972) | 66.7777 | 74.3390 | 46.8826 |
| PB | Private | 13.1228 (+/- 10.4672) | 3.9880 | 11.2795 | 18.8554 |
| PB | Shared | 33.4015 (+/- 14.8031) | 25.2733 | 53.6689 | 51.5773 |
| PB | Residual | 53.4755 (+/- 15.0643) | 70.7389 | 35.0514 | 29.5671 |
| AC | Private | 22.7058 (+/- 16.5534) | 11.6295 | 26.3594 | 50.6683 |
| AC | Shared | 26.9763 (+/- 14.6503) | 44.4749 | 38.9088 | 22.8306 |
| AC | Residual | 50.3177 (+/- 21.2265) | 43.8957 | 35.0514 | 26.5009 |
| STA | Private | 55.1458 (+/- 25.8562) | 48.3895 | 21.5490 | 36.3625 |
| STA | Shared | 23.6595 (+/- 17.2281) | 46.2345 | 66.6929 | 54.3020 |
| STA | Residual | 21.1946 (+/- 21.2975) | 5.3761 | 11.7580 | 9.3353 |

**Table 7. Inhibitory MCFA results**. Values that are reduced compared to bootstrapped aCSF are colored in red and values that are increased compared to aCSF are colored in blue.

DRAFT

| Attribute | Variance type | Variance 1st Trial | Variance 2nd Trial |
|---|---|---|---|
| AP | Private | 11.2873 (+/- 11.81) | 12.4735 (+/- 08.39) |
| AP | Shared | 47.5009 (+/- 15.28) | 38.4021 (+/- 10.44) |
| AP | Residual | 41.2118 (+/- 10.25) | 49.1244 (+/- 08.84) |
| PB | Private | 10.3859 (+/- 08.94) | 19.9091 (+/- 12.32) |
| PB | Shared | 38.0565 (+/- 14.78) | 37.3036 (+/- 14.24) |
| PB | Residual | 51.5576 (+/- 18.05) | 42.7873 (+/- 13.55) |
| AC | Private | 31.0247 (+/- 16.07) | 12.7529 (+/- 09.86) |
| AC | Shared | 35.2139 (+/- 15.73) | 22.0695 (+/- 11.62) |
| AC | Residual | 33.7614 (+/- 14.71) | 65.1776 (+/- 16.40) |
| STA | Private | 21.4966 (+/- 16.40) | 33.4581 (+/- 23.06) |
| STA | Shared | 62.5138 (+/- 17.43) | 48.8704 (+/- 25.72) |
| STA | Residual | 15.9897 (+/- 08.43) | 17.6716 (+/- 15.15) |

**Table 8. Excitatory MCFA results for second aCSF trial**. Values that are reduced compared to bootstrapped aCSF trial 1 are colored in red and values that are increased compared to aCSF trial 1 are colored in blue.

DRAFT

| Attribute | Variance type | Variance 1st Trial | Variance 2nd Trial |
|---|---|---|---|
| AP | Private | 08.4805 (+/- 07.68) | 10.5232 (+/- 07.13) |
| AP | Shared | 56.1049 (+/- 11.24) | 33.9479 (+/- 17.05) |
| AP | Residual | 35.4145 (+/- 07.97) | 55.5289 (+/- 18.73) |
| PB | Private | 08.3097 (+/- 06.94) | 07.4436 (+/- 09.55) |
| PB | Shared | 61.6040 (+/- 12.22) | 42.5329 (+/- 16.00) |
| PB | Residual | 30.0864 (+/- 09.39) | 50.0235 (+/- 15.53) |
| AC | Private | 21.8111 (+/- 10.96) | 23.1401 (+/- 15.18) |
| AC | Shared | 56.9907 (+/- 12.05) | 29.6210 (+/- 10.40) |
| AC | Residual | 21.1982 (+/- 09.24) | 47.2389 (+/- 15.30) |
| STA | Private | 36.8157 (+/- 23.86) | 32.6709 (+/- 21.86) |
| STA | Shared | 37.5418 (+/- 17.49) | 62.5230 (+/- 22.45) |
| STA | Residual | 25.6425 (+/- 22.69) | 04.806 (+/- 02.85) |

**Table 9. Inhibitory MCFA results for second aCSF trial**. Values that are reduced compared to bootstrapped aCSF trial 1 are colored in red and values that are increased compared to aCSF trial 1 are colored in blue.

DRAFT


DRAFT
